# Disentangling Cobionts and Contamination in Long-Read Genomic Data using Sequence Composition

**DOI:** 10.1101/2024.05.30.596622

**Authors:** Claudia C. Weber

## Abstract

The recent acceleration in genome sequencing targeting previously unexplored parts of the tree of life presents computational challenges. Samples collected from the wild often contain sequences from several organisms, including the target, its cobionts, and contaminants. Effective methods are therefore needed to separate sequences. Though advances in sequencing technology make this task easier, it remains difficult to taxonomically assign sequences from eukaryotic taxa that are not well-represented in databases. Therefore, reference-based methods alone are insufficient.

Here, I examine how we can take advantage of differences in sequence composition between organisms to identify symbionts, parasites and contaminants in samples, with minimal reliance on reference data. To this end, I explore data from the Darwin Tree of Life project, including hundreds of high-quality HiFi read sets from insects. Visualising two-dimensional representations of read tetranucleotide composition learned by a Variational Autoencoder can reveal distinct components of a sample. Annotating the embeddings with additional information, such as coding density, estimated coverage, or taxonomic labels allows rapid assessment of the contents of a dataset.

The approach scales to millions of sequences, making it possible to explore unassembled read sets, even for large genomes. Combined with interactive visualisation tools, it allows a large fraction of cobionts reported by reference-based screening to be identified. Crucially, it also facilitates retrieving genomes for which suitable reference data are absent.

## Introduction

Recent advances in sequencing technology are driving largescale reference genome production for a wide range of organisms, especially taxa that have not been sequenced extensively. A key aim of these efforts is to better understand the evolution of these species, along with their roles in ecosystems [1–4]. In addition to the target genome, wild-sourced samples from a species of interest often contain additional genetic material from organelles, cobionts (such as the microbiome, symbionts and parasites), and environmental contaminants. The presence of non-target sequence can be an obstacle to generating a reliable assembly for the target organism, as evidenced by many published genomes that contain contamination [5–8]. Contaminated assemblies can compromise downstream analyses and lead to inaccurate biological conclusions. On the other hand, these data present an opportunity to characterise ecological associations between organisms. Given suitable computational tools, we can also generate reference-quality genomes for cobionts, including unculturable endosymbionts [9, 10].

Efforts such as the Darwin Tree of Life Project (DToL) [2], which aims to sequence 70,000 eukaryotic genomes, provide an unprecedented opportunity to study the evolution of a wide range of organisms and their interacting partners. The high-quality long-read sequencing data being generated as part of the project ought to allow sequences from different sources to be more easily disentangled. In addition to improving assembly contiguity, the reads themselves can be more accurately taxonomically classified [11, 12]. The latter approach is commonly applied to metagenomic analysis of bacterial communities. However, sequences from eukaryotes including animals, plants, fungi and protists remain more challenging to work with than their prokaryotic cousins, which have been sequenced more extensively.

Reference-based methods for sequence binning rely on comparisons to databases, which are often contaminated [5, 8, 13–16], leading to incorrect assignments. In addition, a sufficiently closely related reference may be unavailable for many organisms - especially when considering parts of the tree of life that have not yet been widely explored. The problem becomes even more acute for sequences with high rates of evolutionary divergence. Accordingly, sequences from genomes with a low density of evolutionarily constrained sites, as found in many multicellular organisms, can be difficult to taxonomically assign. Supervised neural network classifiers have similar limitations, as they are, by definition, trained on sequences that have already been discovered. Performance on “out of distribution” samples that do not resemble any data the model has encountered before can therefore be unreliable [17, 18].

How can we reliably separate sequences despite database gaps? Approaches that take advantage of inherent differences in sequence composition between organisms can be helpful. For example, BlobToolKit (BTK) [5] helps to interactively visualise and extract groups of sequences with differing GC content and coverage. Since GC content provides a limited summary of composition and is not always sufficient to distinguish different organisms [19], short substrings (k-mers) can also be used for unsupervised binning. Separating contigs on the basis of k-mer frequencies and coverage, often with the help of dimensionality reduction, is well-established in metagenomics [20–22]. However, the performance of existing tools on mixtures of sequences that include organisms with substantial intragenomic heterogeneity has yet to be explored. Although base composition varies substantially between microbial genomes [23, 24], they are relatively compact and internally homogeneous. Meanwhile, composition in larger genomes is influenced by the spatial distribution of genomic features such as coding sequences, repetitive elements [25, 26], and recombination breakpoints [27–29].

Composition-based clustering of unassembled sequencing reads has also received less attention [30], despite its potential to inform the sequencing and assembly process. The ability to rapidly assess the contents of a read set would provide an opportunity to gauge the quality of a sample prior to assembly - including whether the target genome is present at sufficient coverage. Given the high accuracy of HiFi reads [11], it might be tempting to treat them as short contigs. However, the number of sequences in a read set can run into the millions, creating computational challenges. In addition, differences in sequencing coverage can be helpful in separating different components of a sample [5, 21, 22, 30], but this metric usually depends on the availability of an assembly.

In this work, I examine how visualising annotated low-dimensional representations of sequence k-mer composition can help detect cobionts and contaminants in samples. To address the challenges of working with large read sets, I implemented a Variational Autoencoder (VAE) [31] that projects tetranucleotide counts into two dimensions. VAEs have proven useful for a variety of biological applications, from examining population structure to predicting variant effects and protein function [32–35] Annotating the two-dimensional embeddings learned by the VAE with additional sequence characteristics, such as estimated coding density, further highlights differences in composition between sequences from different sources (Figure 1). I also developed a k-mer based method to approximate coverage, modifying an established procedure for read set profiling [36]. In addition, an interactive dashboard is provided to explore which organisms might be represented in a sample. Rather than attempting to explicitly classify or bin sequences, these tools are intended as part of a multi-layered cobiont identification strategy.

**Fig. 1.**
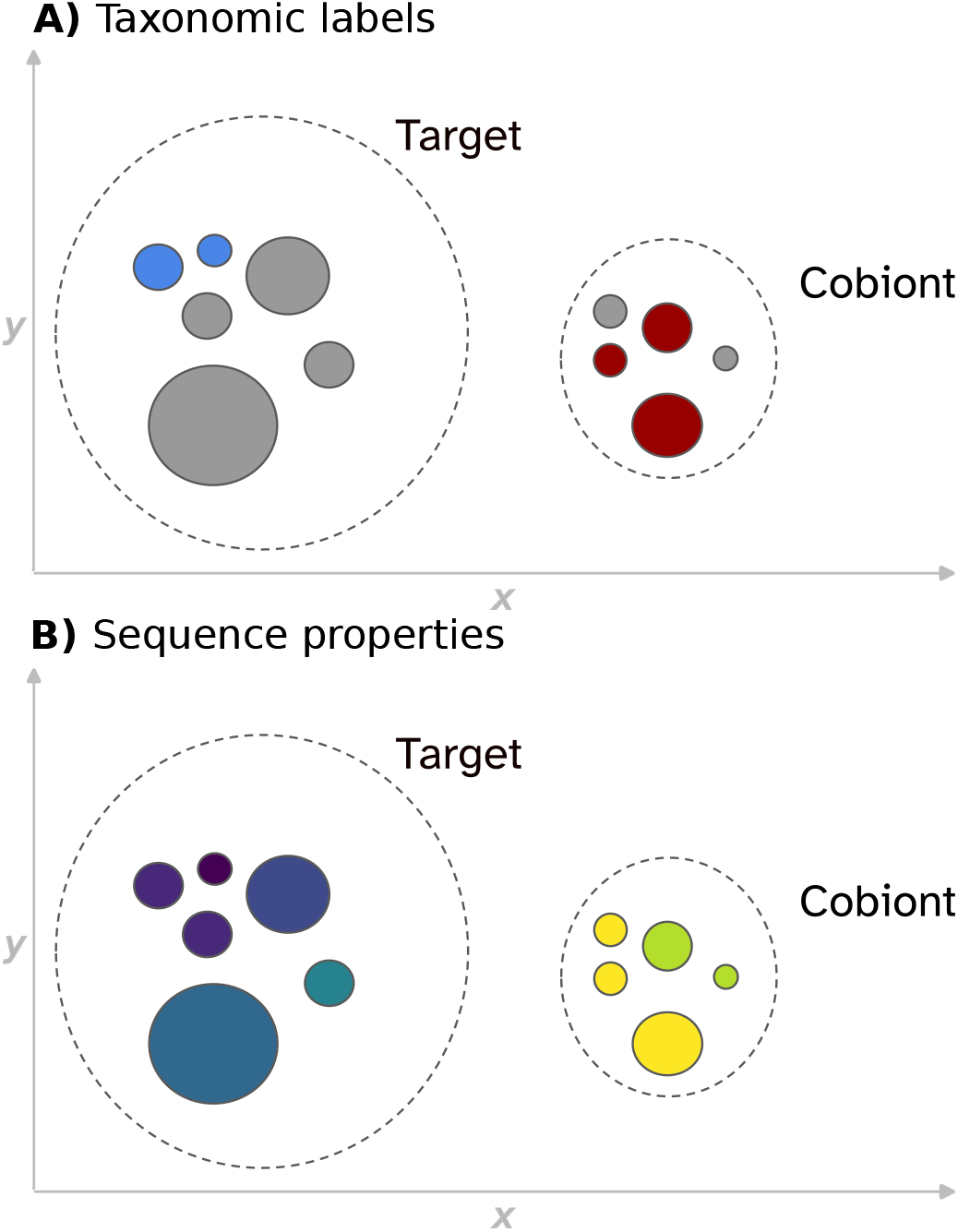
Visualising multiple sources of information about a set of sequences together provides an overview of the components found in the sample. The schematic represents a dataset with sequences from a target species and one cobiont, delineated by dashed lines. Each point represents one sequence, and coordinates reflect two-dimensional representations of tetranucleotide composition learned by a VAE. Sequences with similar composition cluster together. Colours represent sequence annotations. **A)** shows available taxonomic labels. The target is displayed in blue, the cobiont in red, and unassigned sequences in grey. **B)** shows values for sequence statistics (for example, coding density), with lighter colours corresponding to larger values. Where labels are missing in A), information from B) can help differentiate distinct components.

Using data from 204 lepidopterans sequenced by the DToL project [2], I illustrate the impact of taking an integrated approach to cobiont detection and demonstrate that the results are consistent with the output of a reference-based decontamination pipeline. Examples from fish, green algae, and plants show that the composition-based approach can be applied to a broad range of taxa. The VAE allows the workflow to scale to large long-read datasets. In addition, it is able to retrieve cobionts where reference-based approaches commonly fail – particularly where no closely related references are available, as in the case of microsporidians or myxozoans. Finally, I illustrate how target yield can be assessed given a heavily contaminated unassembled read set.

## Methods

### Data

The analyses presented here are based on Pacific Biosciences single-molecule HiFi long reads generated by DToL from single specimens. The 204 lepidopteran species considered, including *Phalera bucephala* and *Blastobasis lacticolella*, are described in [9]. In addition, reads from *Brachiomonas submarina, Viscum album*, and *Thunnus albacares* (from the Vertebrate Genomes Project) were examined. Primary contigs for *P. bucephala*, were assembled with hifiasm version 0.12 using default settings [37]. This assembly is intended for illustration, and does not reflect the released *P. bucephala* assembly.

### Sequence composition

#### K-mer counts

The primary measure of sequence composition considered in this work are canonical tetranucleotide counts (each k-mer and its reverse complement are assigned the same key). In principle, other k-mer sizes *k* could be used, but *k* = 4 provided a reasonable balance between computational cost and the ability to disentangle sequences for a range of samples. Considering the canonical counts reduces the number of features per sequence from 256 to 136 and ensures that reads and their reverse complements cluster together (though note that the full set of 256 captures information about strand asymmetry). To tally counts efficiently, I implemented a Rust program built on the Needletail library. Memory requirements depend on *k* and sequence length, not the number of sequences, making the implementation suitable for datasets with millions of sequencing reads. The k-mer counter is available at https://github.com/CobiontID/kmer-counter.

#### Variational Autoencoders for read decomposition

In order to visualise high-dimensional tetranucleotide count vectors, a VAE is used to reduce the data into two-dimensional space [31, 38]. VAEs consist of a pair of deep neural networks, with an encoder that projects the observed input features (*x*) into a lower-dimensional latent space (*z*), and a decoder that attempts to reconstruct the original input features from samples from the latent space.

Intuitively, sequences with similar tetranucleotide composition will be closer together in latent space than dissimilar sequences. In addition, the decoder’s ability to reconstruct the inputs relies, in part, on the latent space capturing the salient features and structure of the input data. The latent variables can therefore provide information about the processes that generated the observed data [39].

VAEs generally provide better class separation than principal component analysis (PCA), given their ability to learn non-linear projections [32, 40]. This observation holds for read tetranucleotide composition (Figure S7). They are also more computationally efficient than methods such as t-SNE or UMAP [41, 42], making them suitable for large read datasets.

The encoder network of the VAE *q*_*φ*_(*z* | *x*) does not produce deterministic encodings. Instead, for each input and latent dimension, it returns the mean (*µ*) and log variance (log *σ*) of a Gaussian distribution from which random samples are drawn. This makes the model robust to noise in the input data. The decoder *p*_*θ*_(*x* | *z*) then maps the samples back to the original high-dimensional space (see [38], Figure 2.1).

**Fig. 2.**
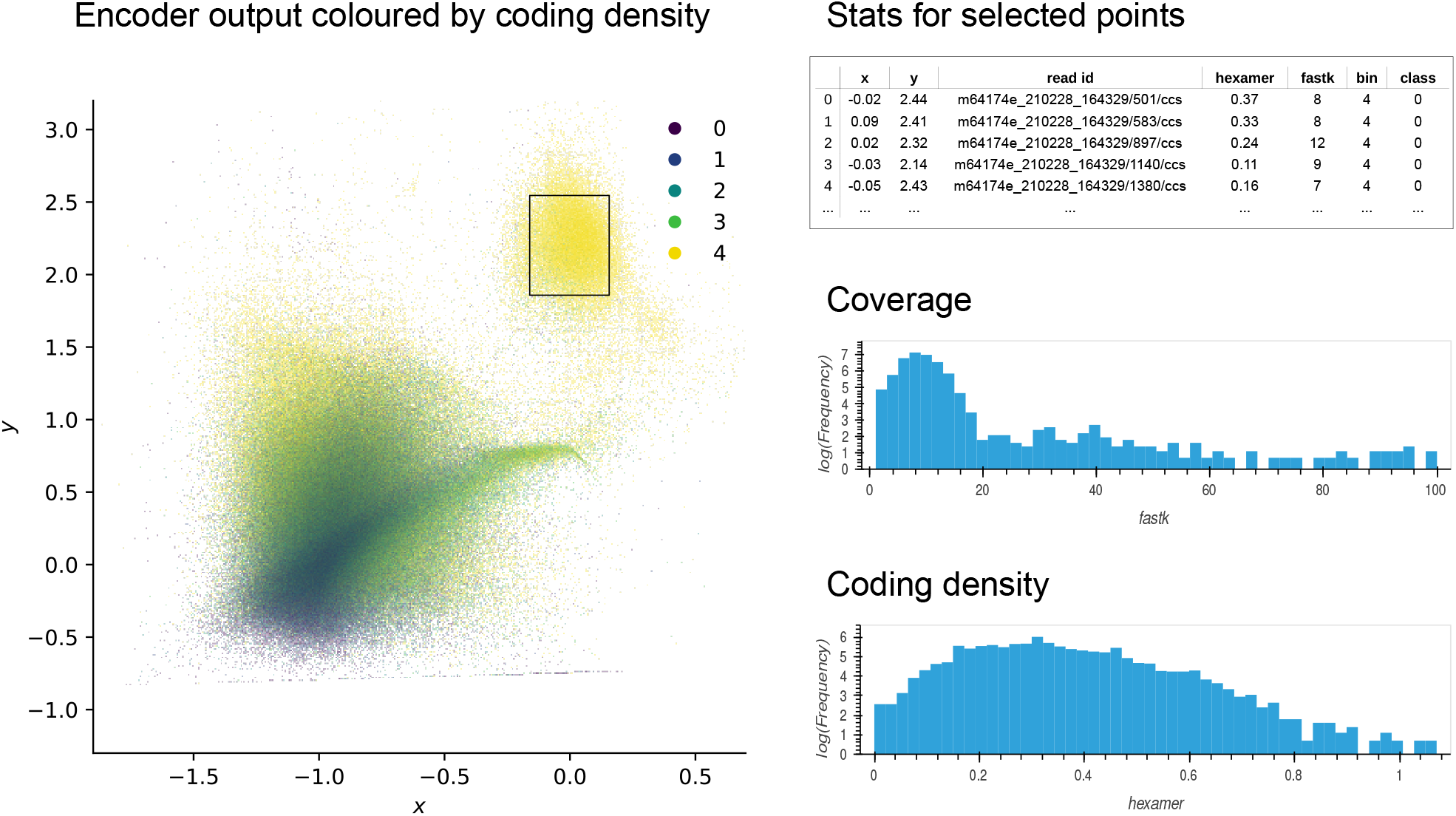
A schematic overview of the interactive dashboard. The left panel displays a colour-coded scatterplot of the 2D coordinates returned by the encoder, which can be filtered by coverage and class (data from *Apamea monoglypha*). The interface permits labelling points by coding density bin, as shown here, or class labels (for example taxonomy). The panels on the right show a summary of the data in the selection defined by the rectangular box. Controls for filtering the data and launching blast queries are not shown here for simplicity.

The variational parameters *φ* and the decoder parameters *θ* are learned by the respective neural networks.

The network is trained by optimising the lower bound on the log likelihood of the data (ELBO) [31], which is given by:

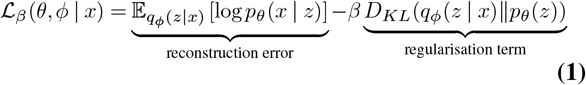

The first term of the model’s loss function serves to minimise the reconstruction error, so that the decoder *p*_*θ*_(*x* | *z*) learns to produce outputs that closely resemble the original input *x*. In addition, a regularisation term encourages the encoder distribution *q*_*φ*_(*z* | *x*) to be close to the unit Gaussian prior *p*_*θ*_(*z*) by minimising the Kullback-Leibler divergence. The latter ensures a compact latent space, where similar inputs are placed close together.

The weighting factor *β* adjusts the penalty on the regularisation term. For the tetranucleotide count data, setting *β <* 1 is necessary to ensure that the model stores useful information about the inputs in the latent space [43], thus allowing sequences to be visually separated (see Figure S6). Hence, the model is a *β*-VAE [44]. Results obtained with other model architectures, such as Associative Compression Networks [40], or VQ-VAE [45], were qualitatively similar. Further details on the implementation of the model and hyperparameter choices are described in the Supplementary Materials. Code is available from https://github.com/CobiontID/read_VAE.

#### Estimated coding density

Given expected differences in coding density between taxonomic groups (low and heterogeneous in most animals, high and relatively homogeneous in bacteria), sequences were additionally annotated with estimated coding densities using hexamer [46], which identifies putative blocks of coding sequence by comparing the composition-controlled 6-mer statistics of windows in a sequence to a reference trained on protein coding sequences. We modified the program to process multi-record fasta files and return the sum of the length of the predicted coding sequence for each record (enabled by setting the *−S* flag) [47]. The estimated coding density is then given by the ratio of the sum of the length of the predicted coding sequences and the total sequence length (values may exceed 1, as both strands are considered). All analyses presented here used the default threshold of 20 and a reference table trained on *Caenorhabditis elegans*.

#### Estimated read coverage

Median k-mer counts across a read set provide an approximation of coverage, provided *k* is sufficiently large that a given substring has a low probability of appearing in a genome multiple times, but small enough to accommodate the sequencing error rate. The median number of times each k-mer in the query sequence appears across the whole set can be extracted from k-mer profiles generated by FastK [36]. Here, *k* = 31, given that compatible tables are routinely produced as part of the DToL quality control pipeline. However, *k* = 60 performed similarly. Note that this approach is not suitable for uncorrected reads with higher error rates, such as uncorrected PacBio CLR or ONT reads. The software used to calculate median counts is available at https://github.com/CobiontID/fastk-medians.

#### Unique k-mers

To help distinguish between reads that belong to DNA molecules that are present in many copies, and reads that contain repetitive sequences, k-mer diversity can be considered in addition to k-mer counts. To calculate this measure, the number of distinct non-canonicalized k-mers in each sequence was counted and divided by sequence length.

To ensure that the maximum possible number of distinct k-mers cannot be smaller than the sequence length, *k* is set so that 4^*k*^ is larger than the expected sequence length of a HiFi read (*k* = 8). The counting software is available at https://github.com/CobiontID/unique-kmer-counts.

### Visualising and exploring reads

Given the large number of reads in each dataset, Datashader [48] was used to render the reduced tetranucleotide data efficiently. Unless otherwise noted, the figures included in this work show the encoder outputs *µ*, which provide sharper cluster boundaries. The x-axis represents the first latent dimension, and the y-axis represents the second latent dimension. To visualise additional sequence statistics such as coding density, continuous values are binned into quantiles and used to colour-code each point. Categorical values, such as taxonomic classifications, can be displayed in a similar manner.

In addition, I provide a Panel [49] dashboard to interactively filter and explore the data (Figure 2). The interface allows the user to zoom in on and select regions of interest, display and download statistics about the selection, inspect sequences, and launch BLAST queries to allow rapid “spot checking” of read clusters (a local server with the reference database loaded into memory ensures fast turnaround times). Rather than explicitly embedding estimated coverage along with the composition feature vector, the interface provides the option of selectively displaying reads within a specified k-mer coverage range. Reads may also be filtered based on class labels.

An interactive demo, complete with an example dataset, is available at https://cobiontid.github.io/reads.html.

### Comparing ascertainment statistics

To examine the circumstances under which sequence composition is helpful in detecting cobionts and contamination, we can consider how often it allowed detection of organisms reported by other tools routinely applied in the DToL assembly and curation process, including NCBI megablast and BTK (see [50], Table 1). In addition, results from MarkerScan were considered [9, 51]. Briefly, MarkerScan uses a profile HMM to search contigs from a sample of interest for small ribosomal subunit (SSU) sequences, and then uses the taxonomic classifications of the SSUs to construct a streamlined Kraken2 database to classify the reads [52]. Candidate cobiont contigs are identified based on whether they are adequately covered by the classified reads, and additional reads mapping to contigs that meet the threshold are identified. Only cobionts with well-covered contigs are recorded, to distinguish them from horizontal transfers.

**Table 1.** Ascertainment of cobionts in 204 lepidopterans. The table gives the fraction of contaminants reported by one of three methods, which could also be detected visually. Each lepidopteran sample was considered separately, and the counts represent the sums across all species. Contaminants that were apparent in the read embeddings but not reported by another pipeline are shown separately in the last column. *Four of the records from curation are due to spurious hits, bringing the true fraction to 6/7 (86%).

|  | Fraction of cobionts retrieved relative to |  |  | Reads only |
| --- | --- | --- | --- | --- |
|  | Mash Screen | Curation | MarkerScan |  |
| All | 137/162 (85%) | 108/135 (80%) | 109/121 (90%) | 116 |
| Bacteria | 136/160 (85%) | 97/111 (87%) | 97/104 (93%) | 94 |
| Fungi | 1/1 (100%) | 6/11 (55%)* | 9/10 (90%) | 12 |
| Metazoa | - | 1/2 (50%) | 0/2 (0%) | 2 |
| Euglenozoa | - | 2/2 (100%) | 2/2 (100%) | 4 |
| Plant | 0/1 (0%) | 2/9 (22%) | 1/3 (33%) | 2 |

For the purposes of this analysis, “detection” was defined in terms of being able to visualise and assign a taxon label by manual spot-checking, and should not be interpreted as classification performance (inconsistency between samples is also expected). In addition to considering static plots annotated with estimating coding density, estimated coverage and the number of unique k-mers per base to identify regions of interest (see Figure S1), coverage filters were applied. Plots were not annotated with taxonomic labels for this analysis.

To automatically identify sequences in different regions of the latent space, reads (*n* = 2) near local peaks in the two-dimensional histogram of the encoder outputs *µ* were randomly sampled, and queried against the NCBI nt database with blast (version 5, 26th June 2021). Peaks were identified by applying the topology method in the *findpeaks* package [53] to an equalised histogram with 200 bins, with a window size of 20.

Where possible, overlaps in the recorded organisms were checked at the family level. However, assignments generated during curation were often above the family level, given that they represent a summary of multiple different screens. Therefore, where the family was not retrievable, the next-highest available level was considered. In order to remove redundant entries, a taxonomic tree was constructed from each set of identifiers, and only nodes with no descendants were retained (hence [*Wolbachia, Rickettsiales*] simplifies to [*Wolbachia*]). The NCBI Taxonomy [54], as implemented in ete3’s NCBITaxa [55], was used to map between levels (database downloaded 3rd October 2022).

For comparisons with Mash Screen [56], a lenient identity threshold of 0.9 was set. Frequent spurious hits to species with extremely low GC content, none of which were confirmed by a second method, were discarded (see Supplementary Material). Since viruses are not amenable to classification using conserved marker genes and are often integrated into the host genome, they are not included in the overall comparison.

## Results

### Sequence composition separates genomes from different sources

To explore how projecting tetranucleotide composition into two dimensions with a VAE allows us to separate different sequence components, I first consider the buff tip moth, *Phalera bucephala* [57], which contains three *Wolbachia* endosymbiont genomes in addition to the target’s nuclear and mitochondrial genomes [9]. Primary contigs belonging to the mitochondrial and *Wolbachia* genomes each form tight clusters that are readily distinguished from the lepidopteran contigs (Figure 3A). In addition to being more homogeneous than the moth sequences, they have notably higher estimated coding density, consistent with a more compact genome architecture.

**Fig. 3.**
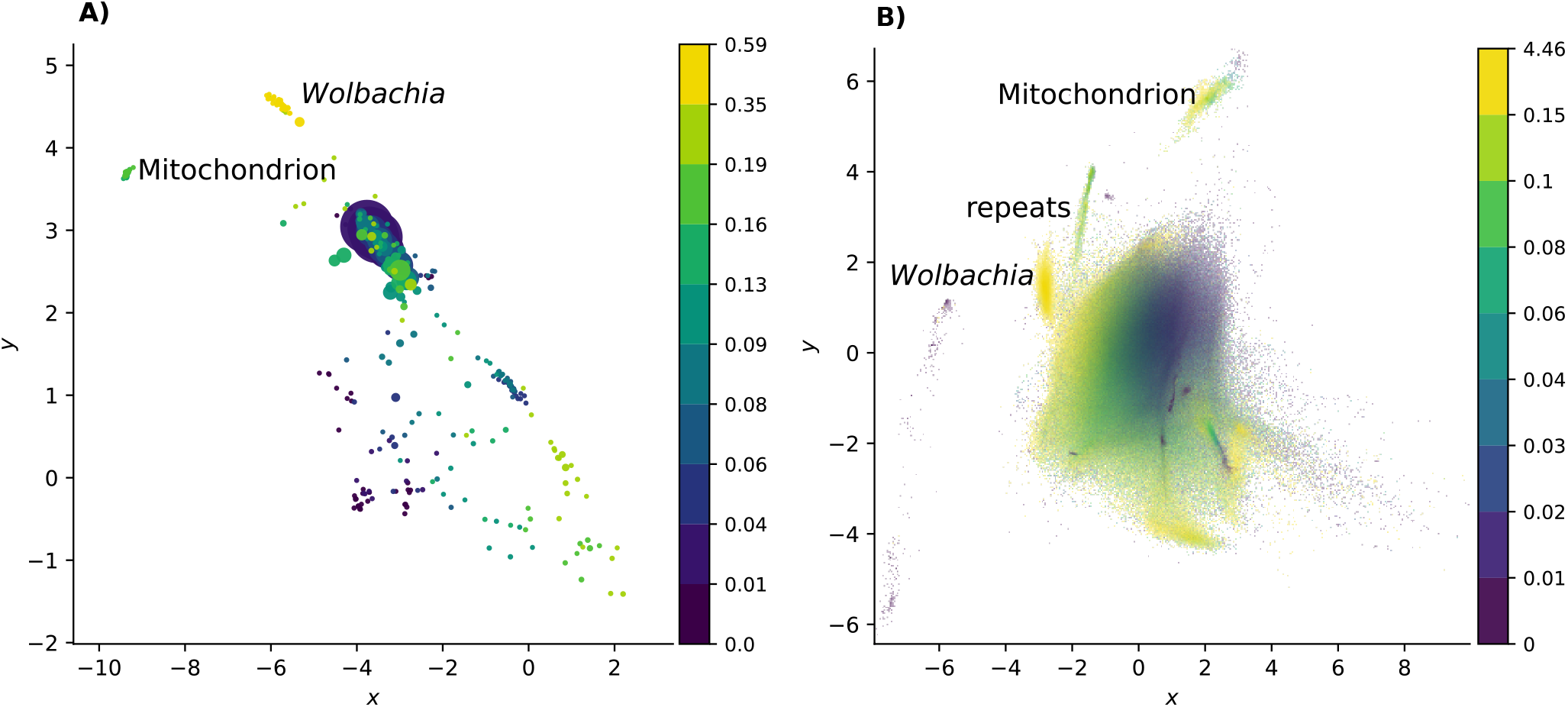
Two-dimensional representations of tetranucleotide count vectors for primary contigs (**A**) and reads (**B**) from the buff tip moth illustrate compositional differences between sequences from different sources. The *x* and *y* axes represent the first and second latent dimensions of a VAE. Each point represents one sequence, and is coloured according to estimated coding density. Each colour represents one decile bin (note that the first two deciles in (B) are combined due to the large number of zeros). Markers in A) are scaled according to contig size. Since contigs and reads were embedded separately, the latent spaces do not align, though the relative relationships between components are similar. The larger moth genome is less coding dense and shows more compositional heterogeneity than the *Wolbachia* genome found in the same sample. The small, homogeneous mitochondrial genome, shows clear separation from the nuclear genome.

Can this approach be extended to unassembled reads? As with the contigs, distinct mitochondrial and bacterial read clusters are apparent (Figure 3B). Read mapping confirms that sequences from the moth nuclear genome, the mitochondrion, and *Wolbachia* are separable in two dimensions (see Figure S1D). The lepidopteran sequences again show more overall heterogeneity, forming multiple clusters.

As expected, *Wolbachia* has overall high coding density, while the moth displays a wider range, reflecting different genomic compartments. In addition, examining the latent representation of the tetranucleotide data highlights a number of groups of repetitive sequences with low k-mer diversity in the moth, including a subset of reads belonging to the heterochromatic female W chromosome [58] (see Figure S1). Hence, decomposed tetranucleotide counts can separate both sequences from different taxa, and sequences from the same genome with different characteristics.

Which features are responsible for separating the different components in the sample? The second latent dimension shows a strong correlation with GC content (*ρ* = *−*0.98, *p <* 0.0001). However, neither dimension alone separates *Wolbachia* and moth sequences (Figure 3B), lending support to the observation that GC is useful but, in some cases, insufficient for identifying contamination. Other samples showed a similar pattern, with GC strongly predicting one latent dimension (see Table S1). The interpretation of the second latent dimension is less clear, with no straightforward alignment with any of the other k-mer statistics considered (see Figures S1, S2, S3, S4).

These results confirm that sequences belonging to the organellar genome and endosymbiont stand out based on their composition and inherent genomic features alone. Examining composition ought, therefore, to be useful for separating sequences even where no suitable reference data are available.

### Retrieval of contaminants ascertained by reference-based approaches

To assess how readily cobionts can be detected from two-dimensional representations of read composition, I examined data from 204 butterflies and moths that had previously been screened for contaminants and cobionts (see [9]). Lepidoptera are a useful test set, given their known associations with bacterial and fungal endosymbionts, and their relatively small genomes. Routine contamination checks included Mash Screen [56] and assessment of scaffolded assemblies with megablast and BTK [5]. In addition, the reads had been classified with MarkerScan [9]. Note that the aim of this analysis was not to assess classification performance, but rather to understand in which settings the approach presented here is likely to be informative.

Overall, composition-based ascertainment of non-lepidopteran sequences via interactive data exploration was high (Table 1). Retrieval of organisms reported by MarkerScan was highest, which is unsurprising as contaminants present at very low coverage are more likely to yield fragmented assemblies with no retrievable SSU region. The sequences of cobionts that were reported by MarkerScan therefore tended to form more conspicuous clusters. Plants and animals, with their heterogeneous genome composition, were difficult to retrieve (but note Figure 5). Bacterial sequences were also overall more easily located, given that their genomes are relatively homogeneous, leading to more compact two-dimensional representations that are discernible even when coverage is low.

Given their compactness, we might expect fungi to behave similarly. At first glance, fungal sequences recorded during the curation process appeared to be surprisingly difficult to retrieve. However, this is likely explained by the absence of a sufficiently closely related reference for fungal sequences found in the moth *Apamea monoglypha*. Though the majority of fungal scaffolds in the sample were assigned to the order *Hypocreales*, consistent with results from MarkerScan and the read data, curation records included four additional orders belonging to the class *Sordariomycetes*. Closer inspection revealed these hits (reported by BTK) to be spurious, and there was no evidence that the corresponding scaffolds were compositionally distinct from those labelled as *Hypocreales* (*A. monoglypha* reads are shown in Figure 2). This alone accounts for 4 of 5 “missing” fungi.

Hence, the apparent inconsistency in performance is not due to limitations of the composition-based approach. In fact, the reference-based approach failed to detect a number of fungal cobionts due to limited reference data. Due to the lack of significant hits, the problem carried over to the BTK screen. MarkerScan and read visualisation were the only approaches consistently able to detect samples infected with nosematids, which are microsporidians, prior to a taxon-specific screen being included in the curation pipeline. As expected, Mash Screen, which relies on exact k-mer matches, performed poorly on non-bacterial cobionts and contaminants. These organisms are less well-represented in reference databases than prokaryotes and often contain a higher fraction of weakly conserved sequence. Of note, Mash Screen rarely detected lepidopteran sequences.

Since manually inspecting each sample may not be feasible, we can also consider an automated approach, sampling reads near local peaks in the two-dimensional histogram of the latent distribution (that is, regions with a higher density of points). However, this resulted in a smaller fraction of organisms being identified, as some contaminants do not form sufficiently prominent peaks. Consistent with this, 76% of previously retrieved MarkerScan records were recovered compared to only 57% of Mash Screen records. In other cases, the sampled sequences had no hits in the nucleotide database, reflecting inherent limitations of the reference-based approach. Drawing additional samples could mitigate the issue, at the expense of runtime. Nevertheless, automatically annotating peaks can highlight a subset of cobiont clusters. Additionally, it can flag sequences that stand out compositionally and cannot be assigned to the target genome.

### Retrieval of cobionts is robust to database gaps

The ability of the composition-based approach to detect microsporidians where other methods do not illustrates the advantages of applying unsupervised learning to cobiont screening – especially in cases where the organism infecting the target belongs to a group of organisms with few available genomic resources. However, as noted in Figure 1, cross-referencing multiple sources of information is a more efficient approach to cobiont identification. To this end, we can annotate tetranucleotide plots with labels generated as part of the MarkerScan pipeline.

In the case of *Blastobasis lacticolella*, reference-based approaches failed to reliably identify reads belonging to *Nosema* due to database gaps. While the SSU is readily classified, visualising the sequences flagged by Kraken2 reveals that around half of the reads in the corresponding cluster remained unassigned (5, 523*/*10, 117). Crucially, mapping the unassigned reads in the cluster against *Nosema* contigs confirms that they in fact belong to the microsporidian (Figure 4). Therefore, combining classification and unsupervised learning can rapidly reveal whether additional undetected cobiont sequences are present.

**Fig. 4.**
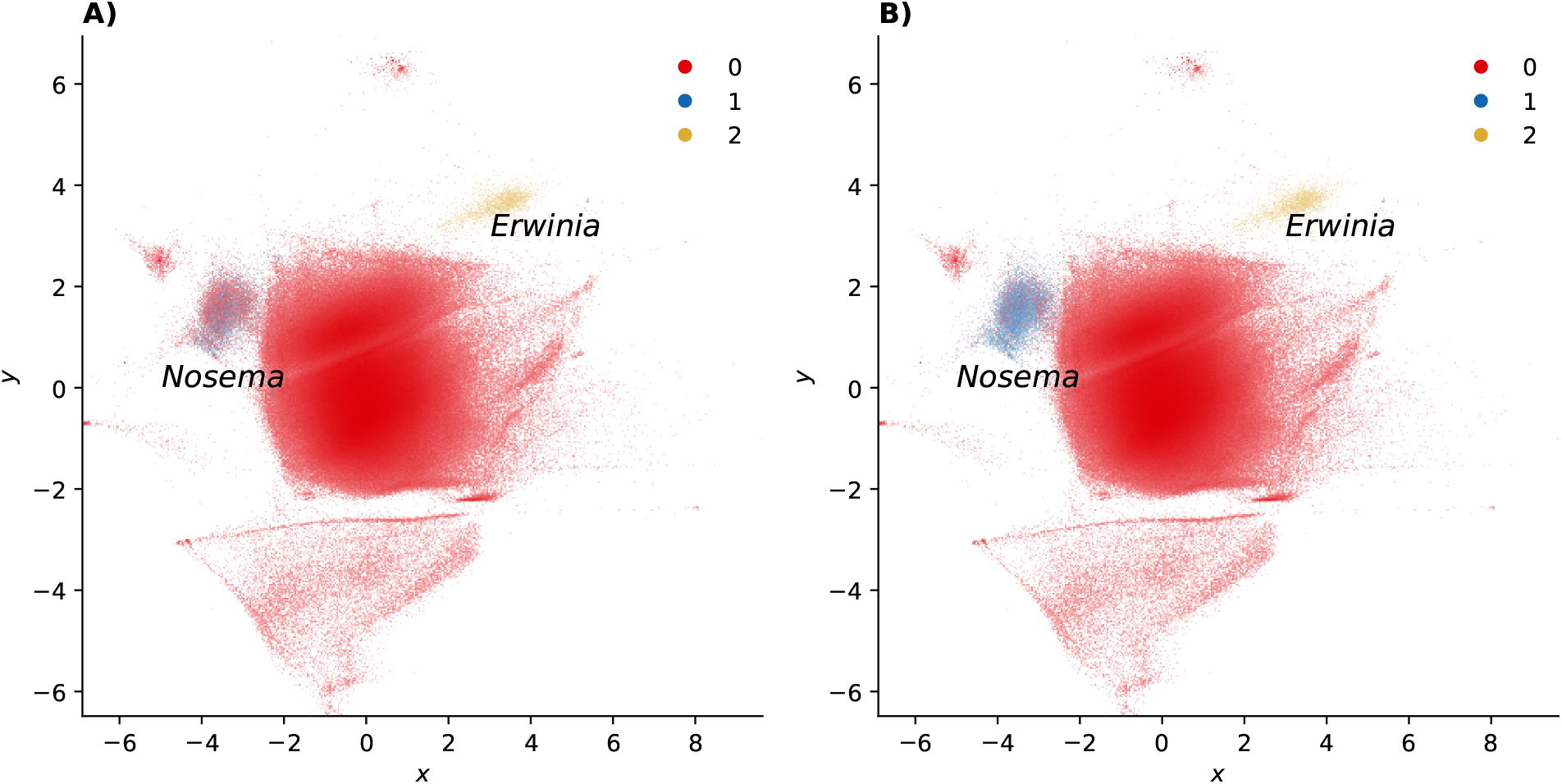
Visualising decomposed reads illustrates that reference-based methods do not retrieve the full set of nosematid sequences present in the London Dowd moth (*Blastobasis lacticolella*). **A)** Reads classified as belonging to *Nosema* (blue) and *Erwinia* (gold) by Kraken2. Unclassified reads are shown in red. **B)** Reads classified by MarkerScan by mapping to possible nosematid contigs, resulting in improved retrieval. VAE batch size was set to 32 to improve visual separation.

**Fig. 5.**
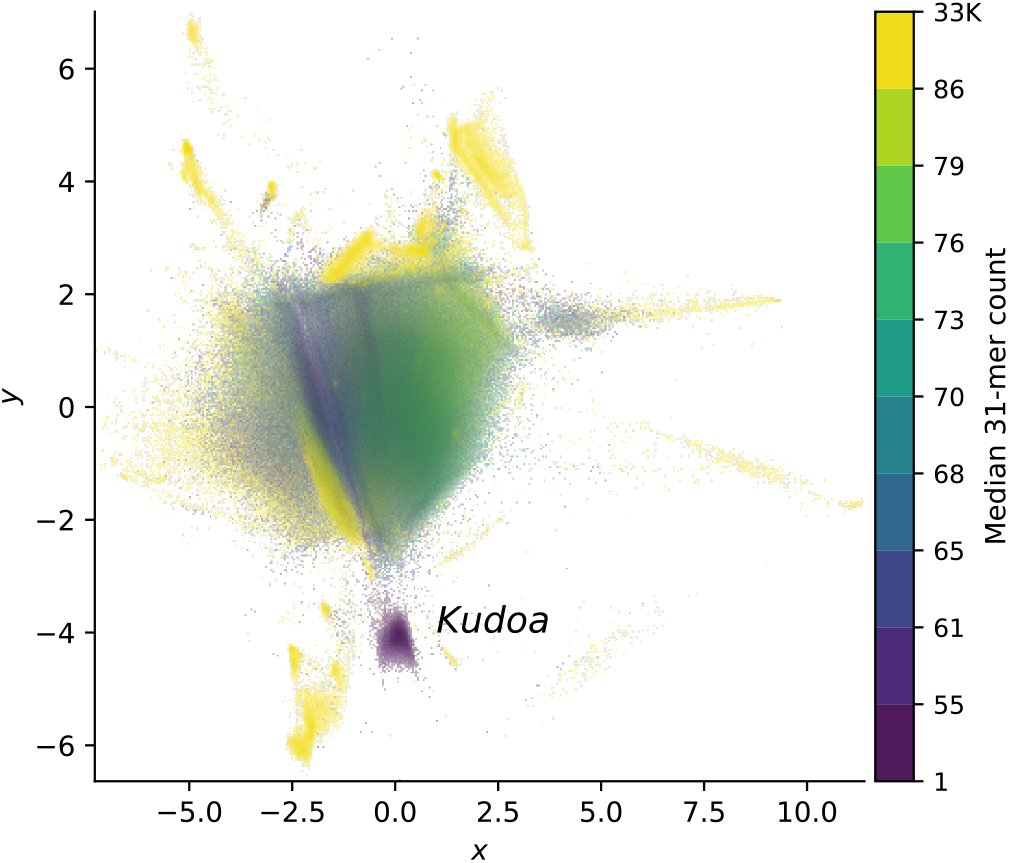
Two-dimensional representations of reads from the yellowfin tuna, *T. albacares*, show evidence of infection with *Kudoa*. Points are coloured by estimated k-mer coverage (*k* = 31), highlighting the difference in coverage between the host and parasite. Repetitive sequences belonging to the host are also apparent.

These observations are not limited to insect-microbe associations. We see a similar pattern in the yellowfin tuna, *Thunnus albacares*, which was infected with a myxosporean parasite of the genus *Kudoa*, a metazoan (manuscript in preparation). Given the large evolutionary distance between the cobiont and the closest relative with an available genome (*K. iwatai* [59]), the fraction of reads classified as *Kudoa* (1, 503) was a mere 5% of those that mapped to *Kudoa* scaffolds removed from the host assembly (30, 542 reads with a minimum mapping score of 50). In fact, the SSU, present on three scaffolds, was key to identifying the parasite. Though reference-based classification of the *Kudoa* reads performed poorly, they form a conspicuous, tight cluster in the tetranucleotide plot, reflecting a highly reduced genome (Figures 5, S3).

### Rapid assessment of unassembled read sets

In both examples discussed above, *Nosema* and *Kudoa* stand out, in part, because their estimated coverage differs markedly from that of the host (see Figures 5, S2). Visualising k-mer coverage as a third dimension through the use of colour can hence highlight non-target sequences. Histograms summarising k-mer coverage are a common tool to assess if a sample has been sequenced to sufficient coverage to assemble successfully. They can also reveal whether significant contamination is present [60, 61]. However, in isolation, they provide limited information about where the target falls in the distribution. Given a histogram for a particular sample, how can we confirm if a peak belongs to the the organism of interest? Cross-referencing two-dimensional representations of read composition and coverage histograms, labelling corresponding coverage ranges with the same colours, is one possible approach.

The green alga *Brachiomonas submarina* provides an illustration. A high peak in the k-mer coverage histogram is apparent at around 2300. However, closer inspection reveals that it is accounted for by sequences from *Brevibacterium linens*, whereas the corresponding peak for the target species is at only around 5-fold coverage (Figure 6). In absence of a preliminary assembly, the identities of the organisms in the clusters can be determined by spot-checking against the NCBI nt database. Sampling reads near local peaks in the two-dimensional distribution confirms that the reads corresponding to the large peak belong to the order *Micrococcales*, with the majority matching the family *Brevibacteriaceae* (see Figure S5). Reads matching the order *Corynebacteriales* are also present in the coverage range overlapping the target’s, but are readily distinguished from the alga based on tetranucleotide composition. Over 85% of reads map to the genome of *Brevibacterium linens strain RS16*, which returned the highest BLAST bitscore for the sampled reads (minimum mapping quality score = 60). Interactively exploring the data leads to similar conclusions.

**Fig. 6.**
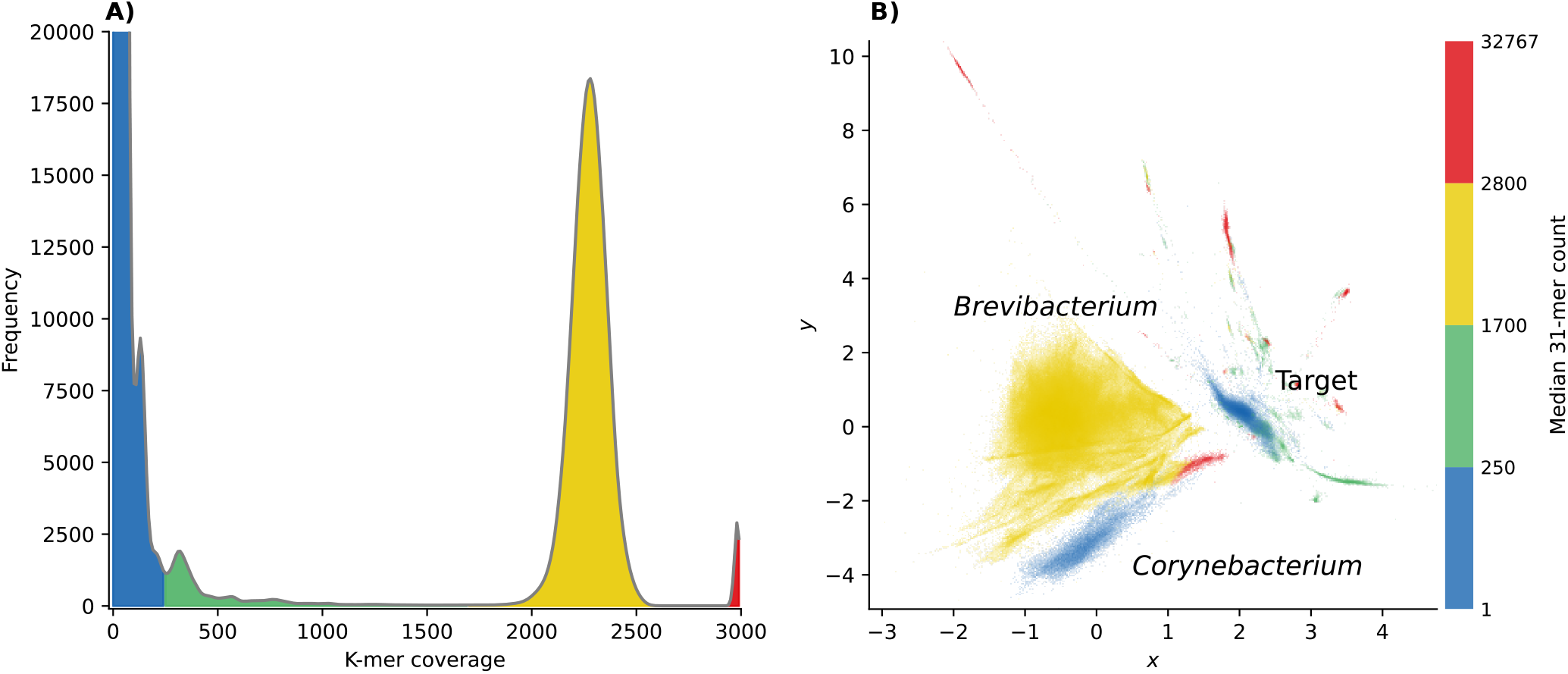
Visualising the k-mer coverage histogram for the green alga *Brachiomonas submarina* alongside read composition allows clusters corresponding to peaks to be identified. The large peak at 2300-fold coverage belongs to *Brevibacterium*, a contaminant. **A)** Count histogram displaying the number of unique k-mers (k = 31) that appear at a given coverage in the dataset. Colours indicate four discrete bands chosen to include conspicuous local peaks in the distribution. **B)** Decomposed read tetranucleotide composition coloured according to k-mer coverage bins shown in A). Note that the distributions of the target reads and *Corynebacterium* overlap, so they cannot be separated based on coverage alone.

Although the coverage ranges in this example were assigned manually for illustration, plots showing automatically assigned bins can be similarly informative. Logarithmic coverage bins indicate that the *Brevibacterium* cluster falls into the same range as the high-coverage histogram peak (Figure S4D). In addition, taken together with the observation that Mash Screen reports two bacterial contaminants, the higher coding density estimates for the cluster hint at the sequences being bacterial rather than algal (Figure S4A). Therefore, automatically generated visualisations of read composition can add context to the more commonly used coverage histograms for rapid sample quality control.

### Runtimes and scalability

The *P. bucephala* read tetranucleotide counts took around 18 minutes to compute on a 2.99GHz CPU (AMD EPYC 7713), with a peak of 5 MB memory usage. Gathering k-mer statistics scales easily to very large read sets. Consider the white mistletoe *Viscum album*, which has 199,755,515 reads: Counting tetranucleotides took 25:42 hours on a single CPU, with a maximum memory usage of 7 MB. Memory usage for estimating the coding density of the mistletoe reads peaked at 8 MB and required 42:01 hours on one CPU (real-world run-times can be reduced by parallel processing).

The VAE required 13:43 minutes and 3951 MB of memory to run for *P. bucephala* (15 epochs, batch size 256). The main computational bottleneck relates to the size of the k-mer count tables. The canonicalised tetranucleotides for *Viscum album* occupy around 100 GB. This translates to an equivalent minimum RAM requirement if the whole array is loaded into memory while training the VAE. In practice, memory usage can be reduced to around 16 GB by loading the preprocessed data in batches, making it feasible to train the model on consumer-grade hardware, albeit at the expense of speed.

## Discussion

This work illustrates how two-dimensional representations of sequence composition can help to identify and separate sequences from different sources in long-read datasets - even where suitable reference sequences are scarce, and the only annotations available are inherent sequence features, such as coding density. As a result, the approach is suitable for a wide range of organisms, including those that are currently not represented in databases. However, it is most effective when integrated with reference-based screens.

In combination with taxonomic labels from other sources, visualising read composition can highlight sequences that belong to an organism of interest but were not identified as such (as with the microsporidian in *B. lacticolella*). It can also flag sequences that are compositionally distinct from reads that belong to known components of the sample, or whose taxonomic assignments appear inconsistent with their sequence features. Identifying gaps and errors in output from classifiers can thus improve ascertainment of cobiont sequences and facilitate assembling complete genomes.

How might these ideas be applied in practice? Given a fragmented cobiont assembly, reads that map to taxonomically labelled contigs or scaffolds may be assembled together with unclassified compositionally similar reads. Composition may also be used to extend the set of candidate cobiont contigs, especially where suitable chromatin conformation data and taxonomic assignments are missing. This is likely to be particularly useful for unculturable organisms with compact genomes, such as microsporidians [62]. Tetranucleotide embeddings are currently being used to help produce more contiguous and complete microsporidian assemblies from DToL samples, as genomes from insect-infecting species are scarce [63]. The approach has also facilitated the assembly of two myxozoan genomes from low-level contamination in two infected fish (manuscript in preparation). In other cases, the tools described here have been used to help distinguish between current infection and horizontal transfer of endosymbiont sequences into the host genome [64–66]. For example, the butterfly *Lysandra bellargus* contains large microsporidian insertions on the W chromosome. In these instances, the structures of the read sets did not resemble infected samples.

As noted in the introduction, the observation that k-mer counts are useful for separating sequences from different microbes is well-established [20, 22]. Indeed, some available binning tools also make use of VAEs [21, 30, 67]. However, the results presented in this work highlight a key consideration for exploring eukaryotic samples. Compositional heterogeneity often results in multiple sequence clusters arising from a single genome, both for reads and assembled sequences (Figure 3). The results of automated binning procedures, like those commonly applied to sequence embeddings in metagenomics, may therefore not be straightforward to interpret. Consistent with this expectation, over-splitting of bins has been reported for mixtures including eukaryotes [51]. A targeted meta-assembly from a selection of reads of interest may therefore be preferable [68], with the assembler performing part of the “clustering” task. This has the advantage of reducing the required computational resources compared to a standard meta-assembly, while taking advantage of information not captured by tetranucleotide counts. In line with this idea, considering connections in assembly graphs improves contig binning [67].

Given the presence of repetitive sequences, coverage estimates are also less obviously useful for separating eukaryotic samples. Both map-based contig coverage and read k-mer multiplicity can deviate from the true number of copies of the genome, though the problem is more pronounced for reads. In addition, in many samples discussed here, prokaryotes that were difficult to separate by composition were present at low coverage with overlapping distributions. Given a setting with only two available latent dimensions, the framework presented here therefore does not explicitly embed estimated coverage, unlike some tools designed for microbes [21, 30, 67]. Instead, coverage is represented using colour, and provided as an interactive filtering criterion. Where separating different strains is of interest, the coverage histogram of a region of interest in the latent space can reveal the number of components present (see Figure 2).

The observations discussed here are based on relatively simple mixtures of sequences: Mostly insect genomes that are overall not heavily contaminated. However, similar principles apply to exploring more complex samples containing many organisms, such as those being catalogued by the Aquatic Symbiosis Genomes Project [3]. The targets often include eukaryotes from groups that have not been extensively sequenced. Clusters of sequences from associated microbes will nevertheless be relatively easy to identify and extract (see Figure S5). In addition, ensuring sufficient target coverage can be challenging for samples with many components. The expected genome size is often unknown. Annotated tetranucleotide embeddings may hence prove useful in identifying the peaks in the coverage distribution corresponding to the genome of interest.

The VAE model used here produces interpretable outputs across a range of long-read datasets (see https://tolqc.cog. sanger.ac.uk for additional examples). However, certain applications and data will benefit from hyperparameter tuning or adjustments to the model architecture [43]. Identifying the optimal settings for a given dataset is notoriously challenging, particularly in an unsupervised setting [32]. Therefore, the specifics of the implementation should be understood as an example, intended to illustrate how two-dimensional representations can help reveal the structure of a mixture of sequences. Performance on contigs and scaffolds is also less consistent than for reads. Other dimensionality reduction methods may therefore be more suitable for high-throughput applications involving relatively small numbers of assembled sequences.

In practice, explicitly integrating taxonomic information and sequence composition might be more effective than a fully self-supervised approach. Where taxonomic labels are available for a subset of sequences, downstream classification performance could be considered as an additional training objective. In this context semi-supervised generative models may prove useful [69, 70]. Rather than attempting to bin sequences after the encoder has been trained, they simultaneously learn to embed and classify data, taking advantage of both labelled and unlabelled inputs. This could provide automated assignments while avoiding the drawbacks of pretrained classifiers. With improved reference-based contamination screens capable of handling diverged sequences [71] and more comprehensive reference data, suitable partially labelled sequences will become more readily available in future.

Another consideration is selecting useful input features. The latent variables of the VAE can be interpreted as factors that give rise to the observed data distribution. Codon models, which parameterise protein-level selection and mutation bias [72–74], are a perhaps more familiar class of generative model describing the evolutionary forces shaping biological sequences. It is not surprising that the organisation of the latent space strongly reflects GC content, consistent with the idea that nucleotide bias is a major driver of genome composition [24, 75]. Though the second latent dimension does not have an obvious interpretation, it contributes to separating sequences. Because tetranucleotide counts provide a simplified and incomplete summary of sequence composition, they are unlikely to capture some salient differences between species. Information about the spatial distribution of sequence patterns is wholly absent. It would therefore be interesting to examine if inputs based on DNA language models offer an advantage.

These analyses demonstrate how 2D representations of sequence composition can be combined with reference-based labels to provide an integrated view of the contents of longread genomic datasets. They also highlight that samples containing eukaryotic genomes require a different approach than prokaryotic metagenomes, given differences in genome structure. The tools presented here are hence intended for exploration, rather than automated binning or classification. VAEs or similar classes of generative model can, however, also provide a framework for identifying sequence features useful for classification. Beyond sample quality control, read sequence embeddings can help retrieve genomes from undersampled taxa – even those not originally targeted.

## DATA AVAILABILITY

The code for the tools presented here is available from https://github.com/CobiontID/ under an MIT license.

## Bibliography

1. Lewin H A, Robinson G E, Kress W J, Baker W J, Coddington J, Crandall K A, Durbin R, Edwards S V, Forest F, Gilbert M T P, et al. 2018, Earth BioGenome Project: Sequencing Life for the Future of Life. Proceedings of the National Academy of Sciences, 115(17):4325–4333.

2. Darwin Tree of Life Project Consortium. 2022, Sequence locally, think globally: The Darwin Tree of Life Project. Proceedings of the National Academy of Sciences, 119(4):e2115642118.

3. McKenna V, Archibald J M, Beinart R, Dawson M N, Hentschel U, Keeling P J, Lopez J V, Martín Durán J M, Petersen J M, Sigwart J D, et al. 2021, The Aquatic Symbiosis Genomics Project: Probing the Evolution of Symbiosis Across the Tree of Life. Wellcome Open Research, 6(254):254.

4. Blaxter M, Archibald J M, Childers A K, Coddington J A, Crandall K A, Di Palma F, Durbin R, Edwards S V, Graves J A, Hackett K J, et al. 2022, Why Sequence All Eukaryotes? Proceedings of the National Academy of Sciences, 119(4):e2115636118.

5. Challis R, Richards E, Rajan J, Cochrane G, and Blaxter M. 2020, BlobToolKit–Interactive Quality Assessment of Genome Assemblies. G3: Genes, Genomes, Genetics, 10(4):1361–1374.

6. Koutsovoulos G, Kumar S, Laetsch D R, Stevens L, Daub J, Conlon C, Maroon H, Thomas F, Aboobaker A A, and Blaxter M. 2016, No Evidence for Extensive Horizontal Gene Transfer in the Genome of the Tardigrade Hypsibius Dujardini. Proceedings of the National Academy of Sciences, 113(18): 5053–5058.

7. Merchant S, Wood D E, and Salzberg S L. 2014, Unexpected Cross-Species Contamination in Genome Sequencing Projects. PeerJ, 2:e675.

8. Francois C M, Durand F, Figuet E, and Galtier N. 2020, Prevalence and Implications of Contamination in Public Genomic Resources: A Case Study of 43 Reference Arthropod Assemblies. G3: Genes, Genomes, Genetics, 10(2): 721–730.

9. Vancaester E and Blaxter M. 2023, Phylogenomic Analysis of Wolbachia Genomes From the Darwin Tree of Life Biodiversity Genomics Project. PLoS Biology, 21(1):e3001972.

10. Kumar S and Blaxter M L. 2011, Simultaneous Genome Sequencing of Symbionts and Their Hosts. Symbiosis, 55(3):119–126.

11. Wenger A M, Peluso P, Rowell W J, Chang P C, Hall R J, Concepcion G T, Ebler J, Fungtammasan A, Kolesnikov A, Olson N D, et al. 2019, Accurate Circular Consensus Long-Read Sequencing Improves Variant Detection and Assembly of a Human Genome. Nature Biotechnology, 37(10):1155–1162.

12. Portik D M, Brown C T, and Pierce Ward N T. 2022, Evaluation of Taxonomic Classification and Profiling Methods for Long-Read Shotgun Metagenomic Sequencing Datasets. BMC Bioinformatics, 23(1):541.

13. Breitwieser F P, Pertea M, Zimin A V, and Salzberg S L. 2019, Human Contamination in Bacterial Genomes Has Created Thousands of Spurious Proteins. Genome Research, 29(6):954–960.

14. Bagheri H, Severin A J, and Rajan H. 2020, Detecting and Correcting Misclassified Sequences in the Large-Scale Public Databases. Bioinformatics, 36(18):4699–4705.

15. Steinegger M and Salzberg S L. 2020, Terminating Contamination: Large-Scale Search Identifies More Than 2,000,000 Contaminated Entries in GenBank. Genome Biology, 21(1):1–12.

16. Orakov A, Fullam A, Coelho L P, Khedkar S, Szklarczyk D, Mende D R, Schmidt T S, and Bork P. 2021, GUNC: Detection of Chimerism and Contamination in Prokaryotic Genomes. Genome Biology, 22(1):1–19.

17. Ponsero A J and Hurwitz B L. 2019, The Promises and Pitfalls of Machine Learning for Detecting Viruses in Aquatic Metagenomes. Frontiers in Microbiology, 10:806.

18. Ren J, Liu P J, Fertig E, Snoek J, Poplin R, Depristo M, Dillon J, and Lakshminarayanan B. 2019, Likelihood Ratios for Out-of-Distribution Detection. Advances in Neural Information Processing Systems, 32.

19. Teeling H, Meyerdierks A, Bauer M, Amann R, and Glöckner F O. 2004, Application of Tetranucleotide Frequencies for the Assignment of Genomic Fragments. Environmental Microbiology, 6(9):938–947.

20. Dick G J, Andersson A F, Baker B J, Simmons S L, Thomas B C, Yelton A P, and Banfield J F. 2009, Community-Wide Analysis of Microbial Genome Sequence Signatures. Genome Biology, 10:1–16.

21. Nissen J N, Johansen J, Allesøe R L, Sønderby C K, Armenteros J J A, Grønbech C H, Jensen L J, Nielsen H B, Petersen T N, Winther O, et al. 2021, Improved Metagenome Binning and Assembly Using Deep Variational Autoencoders. Nature Biotechnology, 39(5):555–560.

22. Alneberg J, Bjarnason B S, De Bruijn I, Schirmer M, Quick J, Ijaz U Z, Lahti L, Loman N J, Andersson A F, and Quince C. 2014, Binning Metagenomic Contigs by Coverage and Composition. Nature Methods, 11(11):1144–1146.

23. Sueoka N. 1962, On the Genetic Basis of Variation and Heterogeneity of DNA Base Composition. Proceedings of the National Academy of Sciences, 48(4): 582–592.

24. Warnecke T, Weber C C, and Hurst L D. 2009, Why There Is More to Protein Evolution Than Protein Function: Splicing, Nucleosomes and Dual-Coding Sequence. Biochemical Society Transactions, 37(4):756–761.

25. Hoyt S J, Storer J M, Hartley G A, Grady P G, Gershman A, de Lima L G, Limouse C, Halabian R, Wojenski L, Rodriguez M, et al. 2022, From Telomere to Telomere: The Transcriptional and Epigenetic State of Human Repeat Elements. Science, 376(6588):eabk3112.

26. Chakraborty M, Chang C H, Khost D E, Vedanayagam J, Adrion J R, Liao Y, Montooth K L, Meiklejohn C D, Larracuente A M, and Emerson J. 2021, Evolution of Genome Structure in the Drosophila simulans Species Complex. Genome research, 31(3):380–396.

27. Weber C C, Boussau B, Romiguier J, Jarvis E D, and Ellegren H. 2014, Evidence for GC-Biased Gene Conversion as a Driver of Between-Lineage Differences in Avian Base Composition. Genome Biology, 15(12):1–16.

28. Galtier N, Piganeau G, Mouchiroud D, and Duret L. 2001, GC-Content Evolution in Mammalian Genomes: The Biased Gene Conversion Hypothesis. Genetics, 159(2):907–911.

29. Galtier N. 2021, Fine-scale Quantification of GC-biased Gene Conversion Intensity in Mammals. Peer Community Journal, 1.

30. Wickramarachchi A and Lin Y. 2022, Binning Long Reads in Metagenomics Datasets Using Composition and Coverage Information. Algorithms for Molecular Biology, 17(1):1–15.

31. Kingma D P and Welling M. 2014, Auto-Encoding Variational Bayes. ICLR.

32. Battey C, Coffing G C, and Kern A D. 2021, Visualizing Population Structure With Variational Autoencoders. G3, 11(1):1–11.

33. Boddé M, Makunin A, Ayala D, Bouafou L, Diabaté A, Ekpo U F, Kientega M, Le Goff G, Makanga B K, Ngangue M F, et al. 2022, High-Resolution Species Assignment of Anopheles Mosquitoes Using K-Mer Distances on Targeted Sequences. eLife, 11:e78775.

34. Frazer J, Notin P, Dias M, Gomez A, Min J K, Brock K, Gal Y, and Marks D S. 2021, Disease Variant Prediction With Deep Generative Models of Evolutionary Data. Nature, 599(7883):91–95.

35. David K T and Halanych K M. 2023, Unsupervised Deep Learning Can Identify Protein Functional Groups from Unaligned Sequences. Genome Biology and Evolution, 15(5):evad084.

36. Fast K. https://github.com/thegenemyers/FASTK.

37. Cheng H, Concepcion G T, Feng X, Zhang H, and Li H. 2021, Haplotype-Resolved De Novo Assembly Using Phased Assembly Graphs With Hifiasm. Nature methods, 18(2):170–175.

38. Kingma D P and Welling M. 2019, An Introduction to Variational Autoencoders. arXiv preprint 1906.02691.

39. Murphy K P. 2023, Probabilistic Machine Learning: Advanced Topics. MIT Press.

40. Graves A, Menick J, and Oord A v d. 2018, Associative Compression Networks for Representation Learning. arXiv preprint 1804.02476.

41. Van der Maaten L and Hinton G. 2008, Visualizing Data using t-SNE. Journal of Machine Learning Research, 9(11).

42. McInnes L, Healy J, Saul N, and Großberger L. 2018, UMAP: Uniform Manifold Approximation and Projection. Journal of Open Source Software, 3(29):861.

43. Wang Y, Blei D, and Cunningham J P. 2021, Posterior Collapse and Latent Variable Non-Identifiability. Advances in Neural Information Processing Systems, 34:5443–5455.

44. Higgins I, Matthey L, Pal A, Burgess C, Glorot X, Botvinick M, Mohamed S, and Lerchner A. 2017, Beta-Vae: Learning Basic Visual Concepts With a Constrained Variational Framework. ICLR.

45. van den Oord A, Vinyals O, and Kavukcuoglu K. 2017, Neural Discrete Representation Learning. CoRR, abs/1711.00937.

46. Durbin R and Thierry Mieg J. The ACEDB Genome Database. In Computational Methods in Genome Research, pages 45–55. Springer, 1994.

47. hexamer. https://github.com/richarddurbin/hexamer.

48. Datashader. https://github.com/holoviz/datashader.

49. Panel. https://github.com/holoviz/panel.

50. Howe K, Chow W, Collins J, Pelan S, Pointon D L, Sims Y, Torrance J, Tracey A, and Wood J. 2021, Significantly Improving the Quality of Genome Assemblies Through Curation. Gigascience, 10(1):giaa153.

51. Vancaester E and Blaxter M. 2024, MarkerScan: Separation and Assembly of Cobionts Sequenced Alongside Target Species in Biodiversity Genomics Projects. Wellcome Open Research, 9(33):33.

52. Wood D E, Lu J, and Langmead B. 2019, Improved Metagenomic Analysis With Kraken 2. Genome Biology, 20:1–13.

53. Taskesen E. findpeaks is for the detection of peaks and valleys in a 1D vector and 2D array (image)., October 2020.

54. Schoch C L, Ciufo S, Domrachev M, Hotton C L, Kannan S, Khovanskaya R, Leipe D, Mcveigh R, O’Neill K, Robbertse B, et al. 2020, NCBI Taxonomy: A Comprehensive Update on Curation, Resources and Tools. Database, 2020: baaa062.

55. Huerta Cepas J, Serra F, and Bork P. 2016, ETE 3: Reconstruction, Analysis, and Visualization of Phylogenomic Data. Molecular Biology and Evolution, 33 (6):1635–1638.

56. Ondov B D, Starrett G J, Sappington A, Kostic A, Koren S, Buck C B, and Phillippy A M. 2019, Mash Screen: High-Throughput Sequence Containment Estimation for Genome Discovery. Genome biology, 20(1):1–13.

57. Boyes D, Holland P W, of Life Consortium D T, et al. 2022, The Genome Sequence of the Buff-Tip, Phalera bucephala (Linnaeus, 1758). Wellcome Open Research, 7(28):28.

58. Sahara K, Yoshido A, and Traut W. 2012, Sex Chromosome Evolution in Moths and Butterflies. Chromosome Research, 20:83–94.

59. Chang E S, Neuhof M, Rubinstein N D, Diamant A, Philippe H, Huchon D, and Cartwright P. 2015, Genomic Insights Into the Evolutionary Origin of Myxozoa Within Cnidaria. Proceedings of the National Academy of Sciences, 112(48): 14912–14917.

60. Vurture G W, Sedlazeck F J, Nattestad M, Underwood C J, Fang H, Gurtowski J, and Schatz M C. 2017, GenomeScope: Fast Reference-Free Genome Profiling From Short Reads. Bioinformatics, 33(14):2202–2204.

61. Ranallo Benavidez T R, Jaron K S, and Schatz M C. 2020, GenomeScope 2.0 and Smudgeplot for Reference-Free Profiling of Polyploid Genomes. Nature Communications, 11(1):1432.

62. Cornman R S, Chen Y P, Schatz M C, Street C, Zhao Y, Desany B, Egholm M, Hutchison S, Pettis J S, Lipkin W I, et al. 2009, Genomic Analyses of the Microsporidian Nosema Ceranae, an Emergent Pathogen of Honey Bees. PLoS Pathogens, 5(6):e1000466.

63. Khalaf A, Francis O, and Blaxter M L. Genome Evolution in Intracellular Parasites: Microsporidia and Apicomplexa. Journal of Eukaryotic Microbiology, page e13033. doi: 10.1111/jeu.13033.

64. Lohse K, Mackintosh A, of Life WI I T, of Life Consortium D T, et al. 2021, The Genome Sequence of the Large White, Pieris Brassicae (Linnaeus, 1758). Wellcome Open Research, 6.

65. Lohse K, Hayward A, Vila R, Howe C, of Life Consortium D T, et al. 2022, The Genome Sequence of the Adonis Blue, Lysandra Bellargus (Rottemburg, 1775). Wellcome Open Research, 7(255):255.

66. Ebdon S, Mackintosh A, Hayward A, Wotton K, of Life Consortium D T, et al. 2021, The Genome Sequence of the Clouded Yellow, Colias crocea (Geoffroy, 1785). Wellcome Open Research, 6(284):284.

67. Lamurias A, Sereika M, Albertsen M, Hose K, and Nielsen T D. 2022, Metagenomic Binning With Assembly Graph Embeddings. Bioinformatics, 38(19): 4481–4487.

68. Feng X, Cheng H, Portik D, and Li H. 2022, Metagenome Assembly of High-Fidelity Long Reads With Hifiasm-Meta. Nature Methods, 19(6):671–674.

69. Makhzani A, Shlens J, Jaitly N, Goodfellow I, and Frey B. 2015, Adversarial Autoencoders. arXiv preprint 1511.05644.

70. Kingma D P, Mohamed S, Jimenez Rezende D, and Welling M. 2014, Semi-Supervised Learning With Deep Generative Models. Advances in Neural Information Processing Systems, 27.

71. Astashyn A, Tvedte E S, Sweeney D, Sapojnikov V, Bouk N, Joukov V, Mozes E, Strope P K, Sylla P M, Wagner L, et al. 2023, Rapid and Sensitive Detection of Genome Contamination at Scale With FCS-GX. bioRxiv, pages 2023–06.

72. Goldman N and Yang Z. 1994, A Codon-Based Model of Nucleotide Substitution for Protein-Coding DNA Sequences. Molecular Biology and Evolution, 11 (5):725–736.

73. Muse S V and Gaut B S. 1994, A Likelihood Approach for Comparing Synonymous and Nonsynonymous Nucleotide Substitution Rates, With Application to the Chloroplast Genome. Molecular Biology and Evolution, 11(5):715–724.

74. Weber C C and Whelan S. 2019, Physicochemical Amino Acid Properties Better Describe Substitution Rates in Large Populations. Molecular Biology and Evolution, 36(4):679–690.

75. Singer G A and Hickey D A. 2000, Nucleotide Bias Causes a Genomewide Bias in the Amino Acid Composition of Proteins. Molecular Biology and Evolution, 17(11):1581–1588.

76. Li H. 2018, Minimap2: Pairwise Alignment for Nucleotide Sequences. Bioinformatics, 34(18):3094–3100.

77. Loaiza Ganem G and Cunningham J P. 2019, The Continuous Bernoulli: Fixing a Pervasive Error in Variational Autoencoders. Advances in Neural Information Processing Systems, 32.

78. Kingma D P and Ba J. 2014, Adam: A Method for Stochastic Optimization. ICLR).

79. Alemi A, Poole B, Fischer I, Dillon J, Saurous R A, and Murphy K. 2018, Fixing a broken ELBO. In International Conference on Machine Learning, pages 159–168. PMLR.

80. Menon S, Blei D, and Vondrick C. 2022, Forget-Me-Not! Contrastive Critics for Mitigating Posterior Collapse. In Uncertainty in Artificial Intelligence, pages 1360–1370. PMLR.

81. Hoffman M D and Johnson M J. 2016, ELBO Surgery: Yet Another Way to Carve Up the Variational Evidence Lower Bound. In Workshop in Advances in Approximate Bayesian Inference, Neural Information Processing Systems, volume 1.

82. Asperti A and Trentin M. 2020, Balancing Reconstruction Error and Kullback-Leibler Divergence in Variational Autoencoders. IEEE Access, 8:199440–199448.

