## Supplementary Material for "Disentangling Cobionts and Contamination in Long-Read Genomic Data using Sequence Composition"

### Read plot annotations

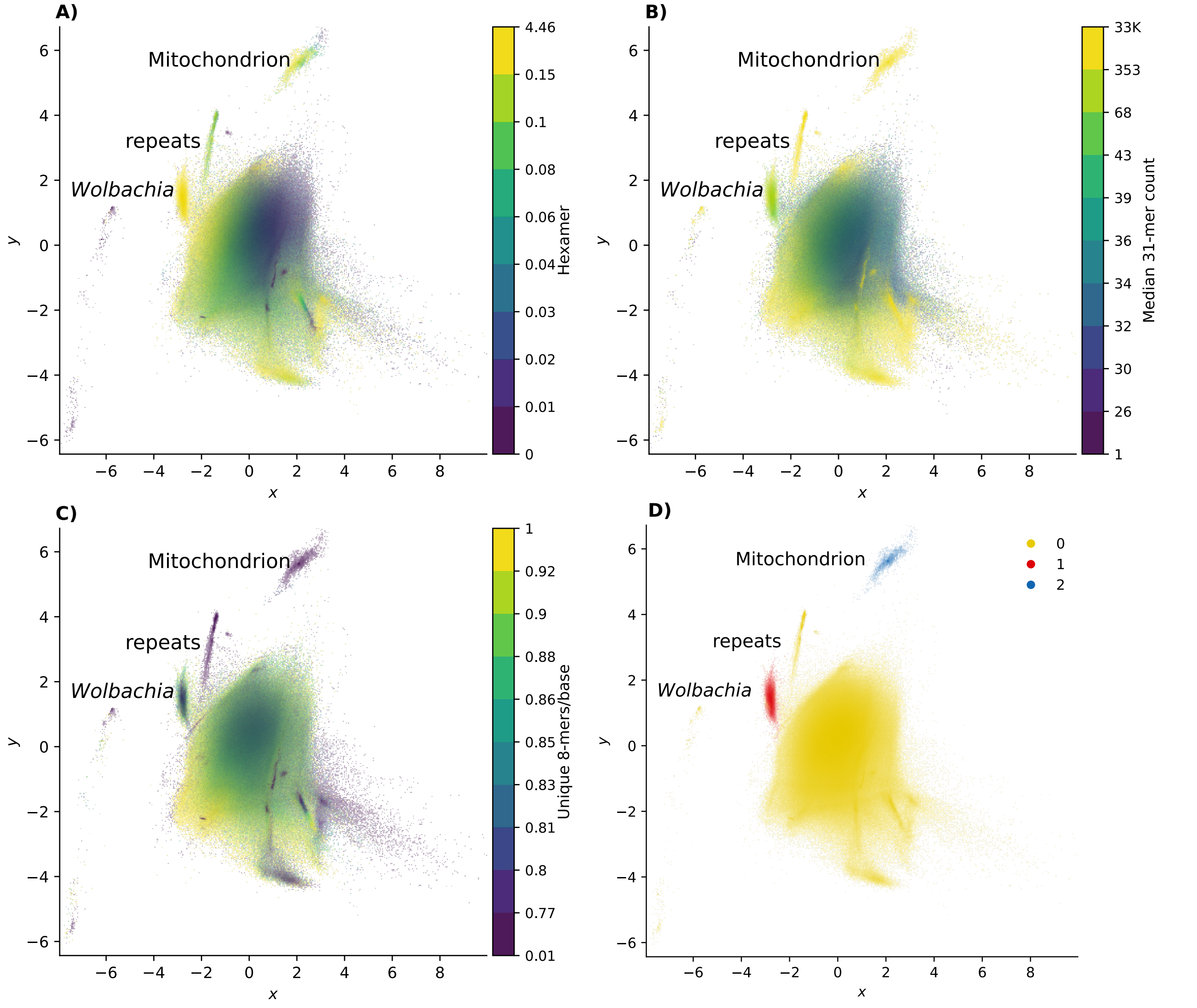


**Supplementary Figure 1.** Decomposed tetranucleotide counts for reads from *Phalera bucephala*, annotated according to **A)** estimated coding density; **B)** estimated k-mer coverage ($k = 31$); and **C)** k-mer diversity, measured by the number of distinct k-mers per base ($k = 8$). Low scores indicate sequences that tend to repeat k-mers, accounting for some clusters with high k-mer coverage estimates. The W-linked sequences above the *Wolbachia* cluster, labelled "repeats" in the plots, represent one example of this. **D)** Read mapping with minimap2 [76] confirms that the VAE model separates sequences from different sources. Reads are colour-coded based on whether they map to the *P. bucephala* nuclear genome (yellow), the *P. bucephala* mitochondrial genome (blue; GenBank: LR990640.1), or one of the three complete *Wolbachia* genomes extracted from the sample (red) [9] with a score greater than zero. Horizontally transferred sequences, such as NUMTs, group with compositionally similar sequences.

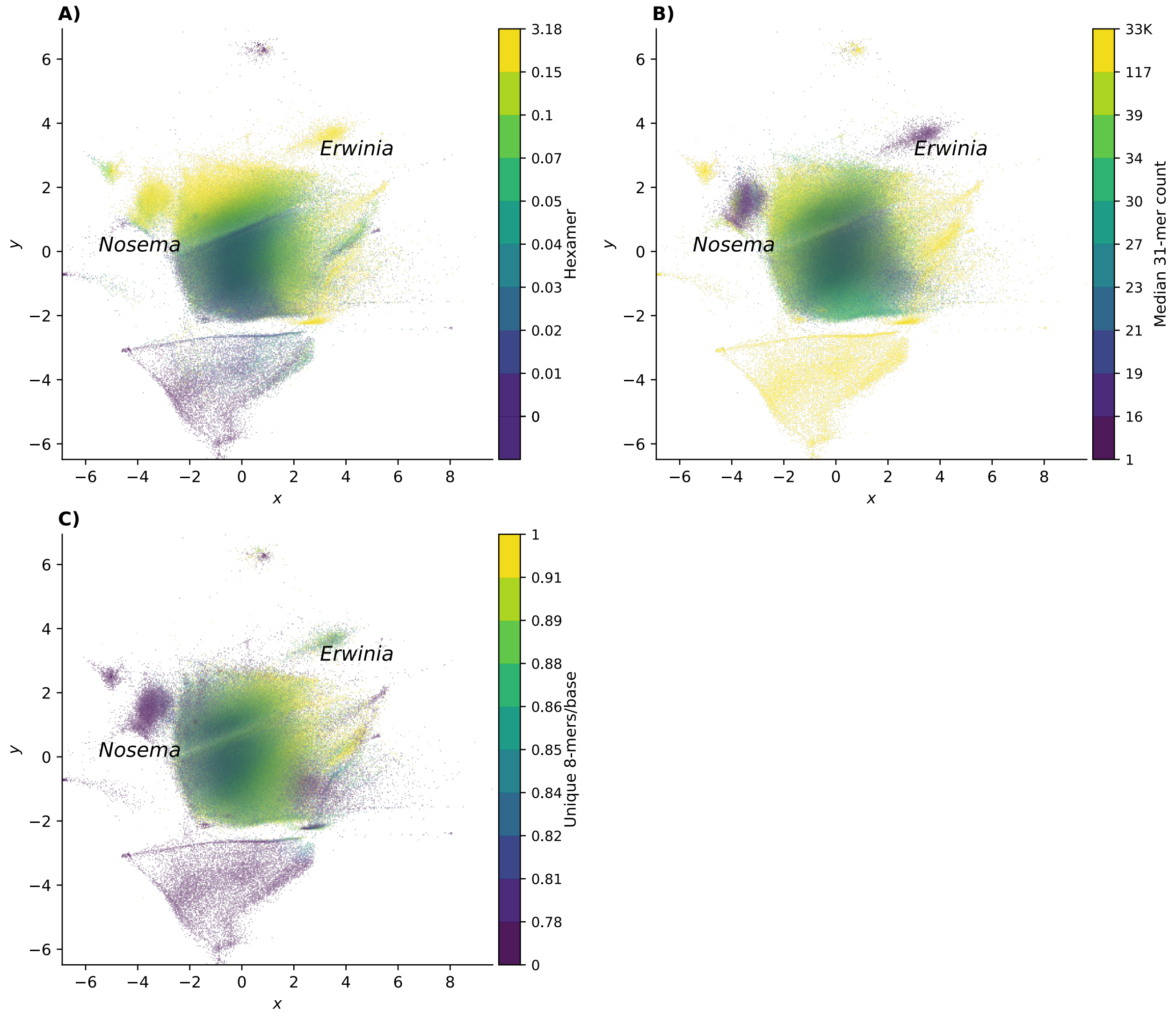


**Supplementary Figure 2.** Annotated read plots for *Blastobasis lacticolella*. *Erwinia* and *Nosema* sequences have high coding density and low coverage. *Nosema* sequences have low k-mer diversity, consistent with high genomic AT-content. Note the presence of repeats belonging to the moth with high median k-mer coverage. **A)** Estimated coding density **B)** Estimated k-mer coverage $(k = 31)$ **C)** K-mer diversity (number of distinct k-mers/base for $k = 8$).

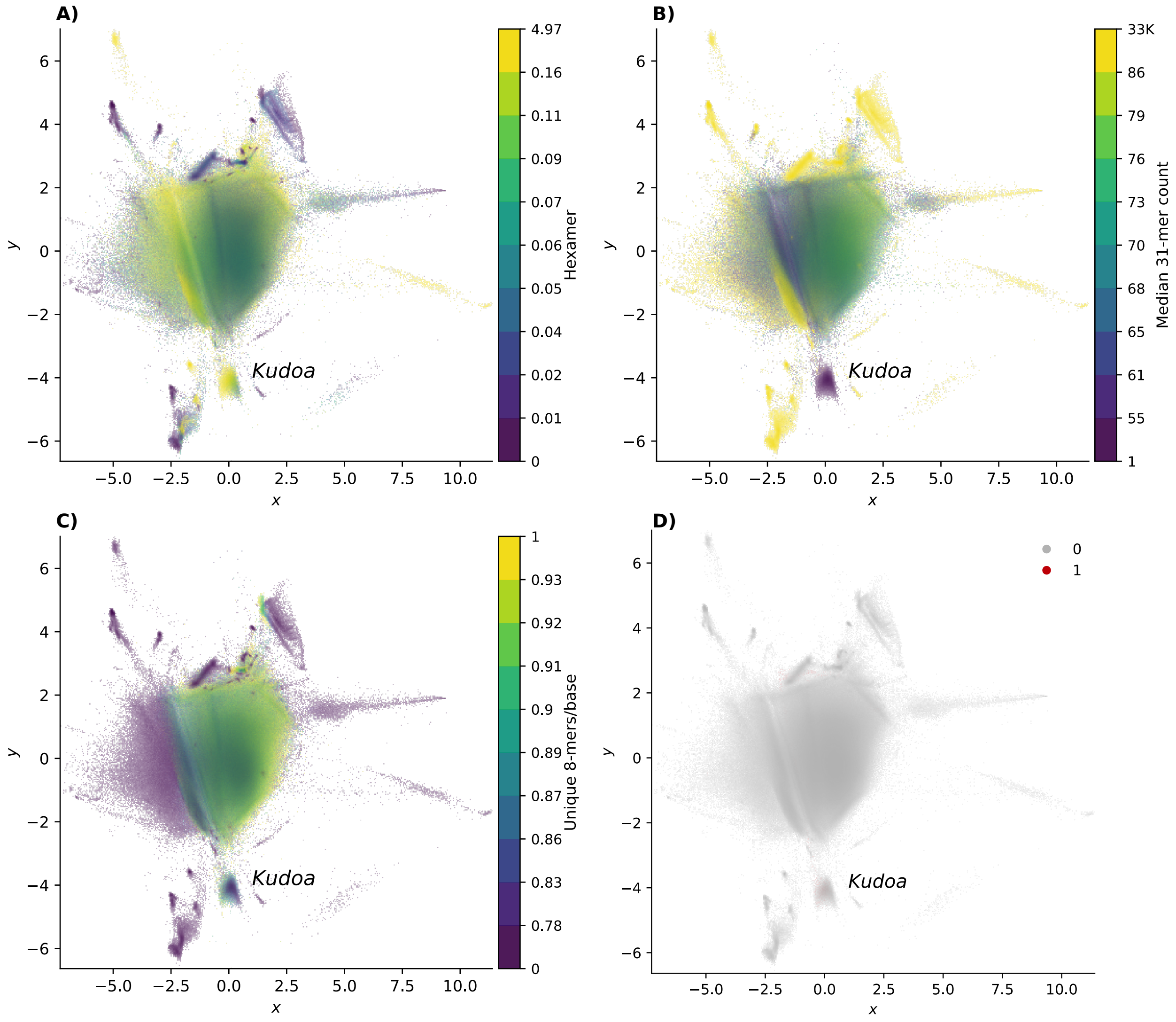


**Supplementary Figure 3.** Annotated read plots for the yellowfin tuna, *T. albacares*. **A)** Estimated coding density **B)** Estimated k-mer coverage $(k = 31)$ **C)** K-mer diversity (number of distinct k-mers/base for $k = 8$) **D)** Reads identified as belonging to *Kudoa* by Kraken 2 marked in colour, with the remaining reads marked in grey. The coloured reads are almost imperceptible, highlighting the high false-negative rate.

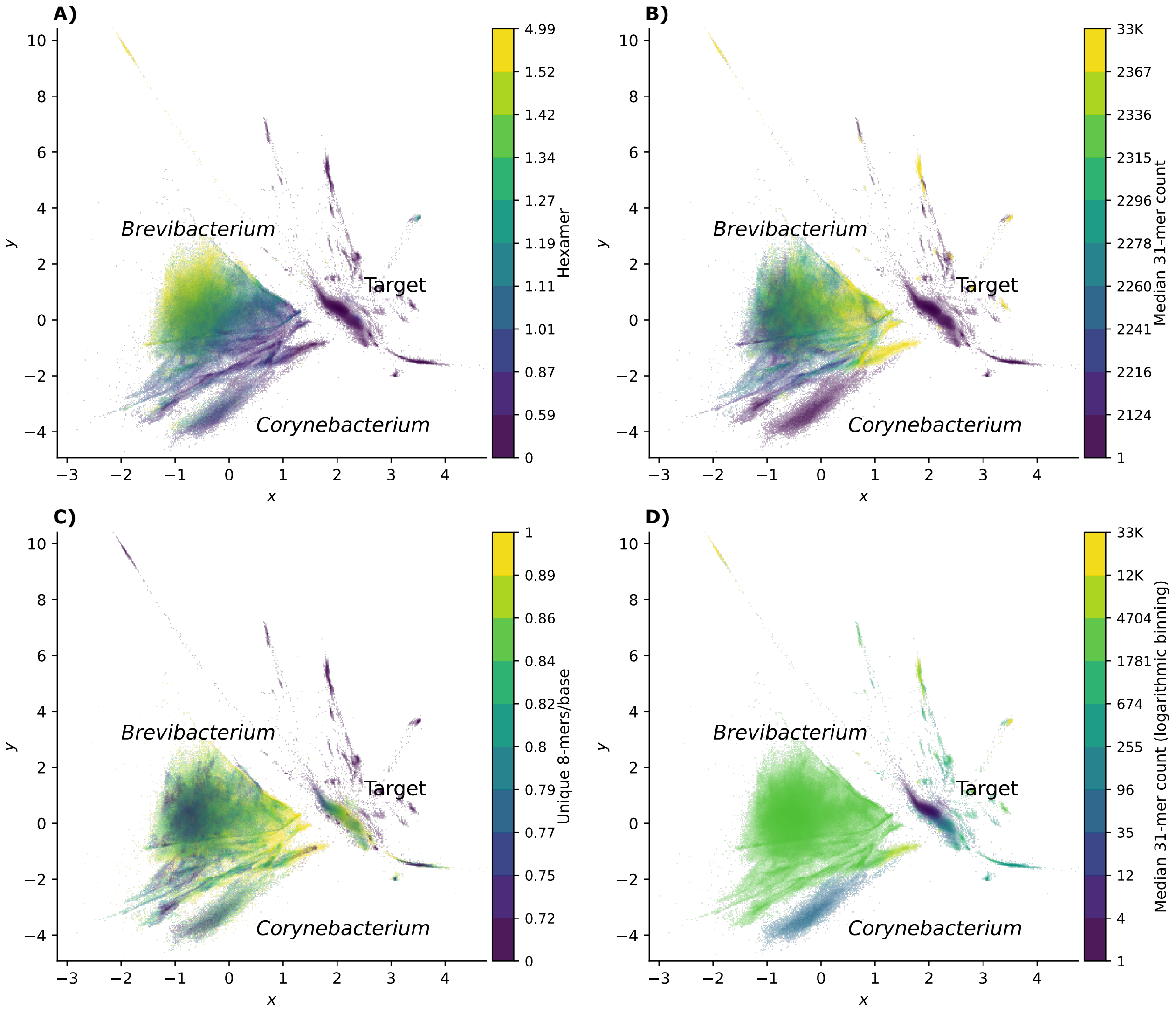


**Supplementary Figure 4.** Annotated read plots for *Brachiomonas submarina*. **A)** Estimated coding density **B)** Estimated k-mer coverage ($k = 31$) **C)** K-mer diversity (number of distinct k-mers/base for $k = 8$). Panels **A**-**C** show decile bins. **D)** Estimated k-mer coverage with logarithmic bins ($k = 31$). Given the k-mer coverage distribution of this sample (Figure 6) discretizing the values using logarithmic equal-width binning provides a more informative overview of the approximate coverage of each component of the mixture.

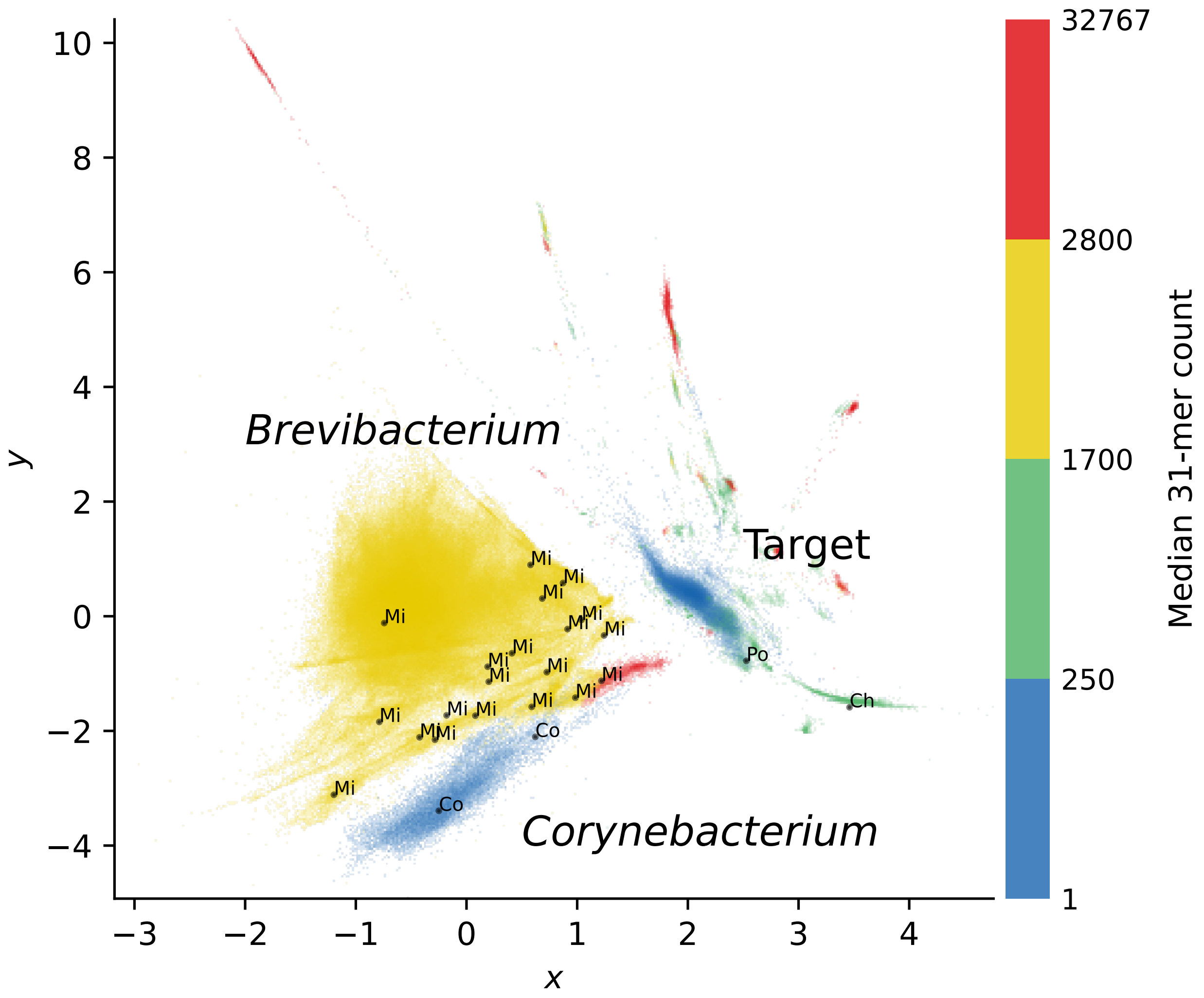


**Supplementary Figure 5.** Annotated read plot for *Brachiomonas submarina* showing locations of sampled reads and corresponding nucleotide blast results, labelled by taxonomic order (bins = 100). For each read, the hit with the highest bit score is considered. To simplify the plot, only the first read with a hit is shown for each sampled peak, though results are consistent across all reads. “Mi” denotes hits to *Micrococcales*, “Co” to *Corynebacteriales*, “Ch” to *Chlamydomonadales* (chloroplast), and “Po” to *Poales*. The latter hit is spurious, with a low bit score (178), although it correctly assigns the sequence to *Viridiplantae*, which illustrates the difficulties identifying algal sequences using sparse reference data. The colours represent bands in the k-mer coverage histogram, as in Figure 6.

### VAE implementation and choice of hyper-parameters

The VAE model presented here is implemented using Keras/TensorFlow2 with a custom GradientTape training loop, based on code adapted from https://keras.io/examples/generative/vae/. To account for variation in sequence length, each tetranucleotide feature vector is divided by its sum, and min-max scaling is then applied to each column to give a range between 0 and 1. These scaled vectors are the input $X$ passed to the neural network.

#### *Loss function*

As described above, the model objective is given by:

$$\mathcal{L}_\beta(\theta, \phi \mid x) = \underbrace{\mathbb{E}_{q_\phi(z|x)}[\log p_\theta(x \mid z)]}_{\text{reconstruction error}} - \beta \underbrace{D_{KL}(q_\phi(z \mid x) \| p_\theta(z))}_{\text{regularisation term}}$$

To compute the reconstruction error term (left), a continuous Bernoulli correction [77] is applied to the binary cross-entropy loss of the reconstruction, to accommodate the bounded distribution resulting from min-max scaling of the length-normalised k-mer count data. The choice of the parameter $\beta$ that determines the weight given to the regularisation term (right) is described in more detail below (“Mitigating posterior collapse").

#### *Architecture and training*

To optimise the model, the encoder and decoder networks, which each contain three hidden layers, are trained end-to-end using the Adam optimiser [78] for up to 15 epochs with a default mini-batch size of 256, an initial learning rate of 0.001, and a validation split of 0.2. After the first fully connected layer of the encoder network, dropout (p = 0.2) and batch normalisation are applied. This is mirrored in the decoder. The hidden fully connected layers all use a rectified linear unit (ReLu) activation function. The sigmoid activation applied to the decoder's final layer gives a range of outputs between 0 and 1. If the validation loss fails to improve for three epochs, the learning rate is scaled by a factor of 0.2, down to a minimum of 0.00001. In addition, training stops when the validation loss fails to improve for four epochs. The random seed may be fixed, to allow for easier comparison between different hyperparameter settings for a given set of sequences. Plots of the latent embeddings at the start of training and after each epoch can optionally be saved in order to visually track training.

Selecting an optimal set of hyperparameters for a VAE is known to be challenging [32]. Performing an exhaustive grid search for hundreds of read sets is not feasible, particularly given that the model is not supervised - that is, no ground truth is available to evaluate performance against. Because suboptimal solutions that do not capture the underlying structure of the data well can result in a small ELBO [79], identifying the settings that result in the smallest loss is insufficient.

#### *Mitigating posterior collapse*

A weakness of the VAE is that it is prone to posterior collapse, where one or more latent variables become “inactive", and the encoder $q_\phi(z \mid x)$ simply reproduces the prior $p(z)$ (see [39]). The model fails to store information that is useful for reconstruction in the latent codes, and therefore $z$ does not capture the underlying structure of the data well. In practice, this results in poor visual separation of dissimilar sequences (Figure S6). One interpretation of this problem is that the KL regularisation term discourages mutual information between the input $x$ and latent representations $z$ [80, 81], and $x$ and $z$ therefore become independent.

Adjusting the weight $\beta$ assigned to the term can help avoid parameters that lead to latent collapse [43]. While setting $\beta > 1$ theoretically encourages the model to learn “disentangled” representations, where the latent dimensions each represent a different aspect of the variation found in the input data, it also encourages the model not to use the latent codes. On the other hand, when $\beta$ « 1, more information tends to be stored in $z$ [39]. For the above model and the data considered in this work, setting $\beta = 0.0025$ provides a reasonable balance, with good separation between different sample components, as illustrated in Figure S6. This is coincidentally equivalent to the scaling factor used by VAMB [21] for two latent dimensions, while LRBinner [30] uses a weight of 0.002.

For some samples, the defaults above nevertheless lead to latent collapse. This is not surprising, as posterior collapse is a function of both the dataset and the model [43]. To mitigate this problem, the implementation described here provides the option to track the mean variance of each latent variable during training [82]. Values close to one indicate that the latent variable is not informative, and can be set to automatically trigger a reduction in $\beta$ at the end of the epoch. In many cases, this will “rescue” training and result in a plot that provides at least some insight into the structure of the dataset. Although the callback is not likely to identify the optimal value for $\beta$, it provides a mechanism to programmatically identify datasets where the defaults require tuning.

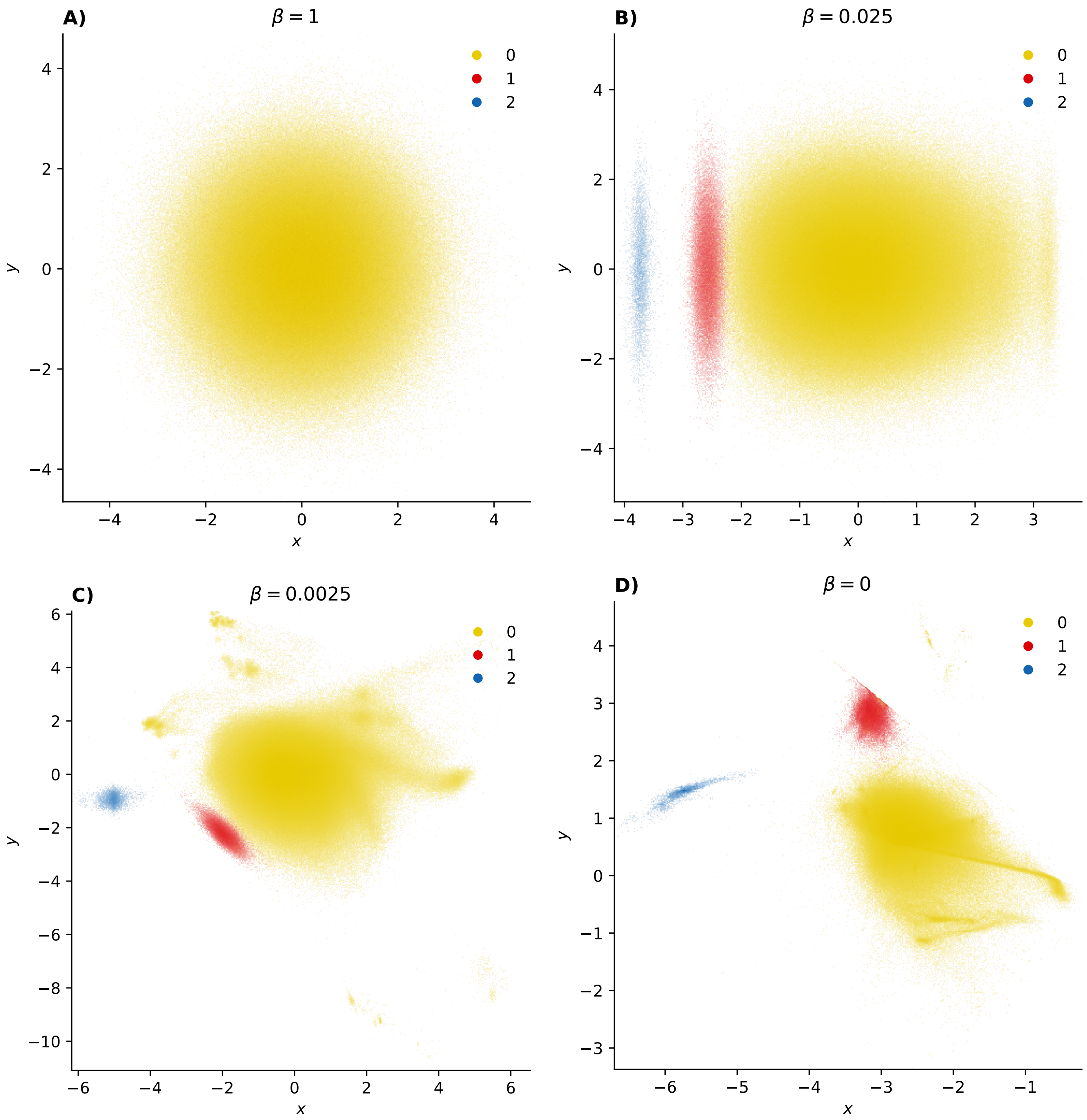


**Supplementary Figure 6.** The influence of the weight placed on the regularization loss on the latent codes $z$ is illustrated with reads from the buff tip moth (*P. bucephala*). Two-dimensional representations of reads are colour-coded based on whether they map to the moth nuclear genome (yellow), the moth mitochondrion (blue), or *Wolbachia* (red), as in Figure S 1D. Note that the examples here show $z$ rather than $\mu$, as $z$ better reflects the uncertainty in the latent distribution. **A)** Latent samples generated with $\beta = 1$ (corresponding to the standard VAE) show no separation between groups, as $q_\phi(z|x)$ has collapsed to the prior $p(z)$. **B)** With $\beta = 0.025$, some separation between reads from different sources is apparent along the first latent dimension (x-axis), but fine-scale structure within the moth nuclear genome is not discernible and the second dimension has collapsed. **C)** Samples generated with $\beta = 0.0025$ show separation between sequences from different components of the sample along both latent dimensions, and heterogeneity within the moth nuclear genome is captured. **D)** When $\beta = 0$ (no regularisation), the model imposes no explicit constraints on the latent space. As a result, the data are not centred around zero, and the latent representations are less compact.

#### ***Embedding coverage information***

Some methods (e.g. VAMB [21]) take coverage estimates as input and structure the model so that the decoder reconstructs composition and coverage separately. However, tests on insect data showed no clear practical improvement in sequence separation for two latent dimensions. Here, the base model was extended to embed the natural logarithm of the median k-mer coverage for each read, applying a mean squared error loss to the reconstruction. The coverage vector tended to “take over” one of the two latent dimensions, leaving a single dimension to capture tetranucleotide composition. A very slight advantage could perhaps be noted for small batch sizes (e.g. 16) in some cases. However, reducing the batch size substantially increases the time required to train the model. The “tetranucleotide + coverage” model is, therefore, less suited to high-throughput screening of read datasets that include eukaryotes (see Discussion for limitations of coverage estimates from heterogeneous genomes).

### Comparison with PCA

As expected, projecting read tetranucleotide count data into two dimensions using principal component analysis provides some separation between sequences from different sources, but the boundaries tend to be less distinct compared to the latent embeddings from a VAE (see Figure S7). UMAP and t-SNE were too computationally expensive to apply to large read sets, and are therefore not considered here. Therefore, although PCA is fast and can provide a quick snapshot for particularly large datasets, it is insufficient in some cases.

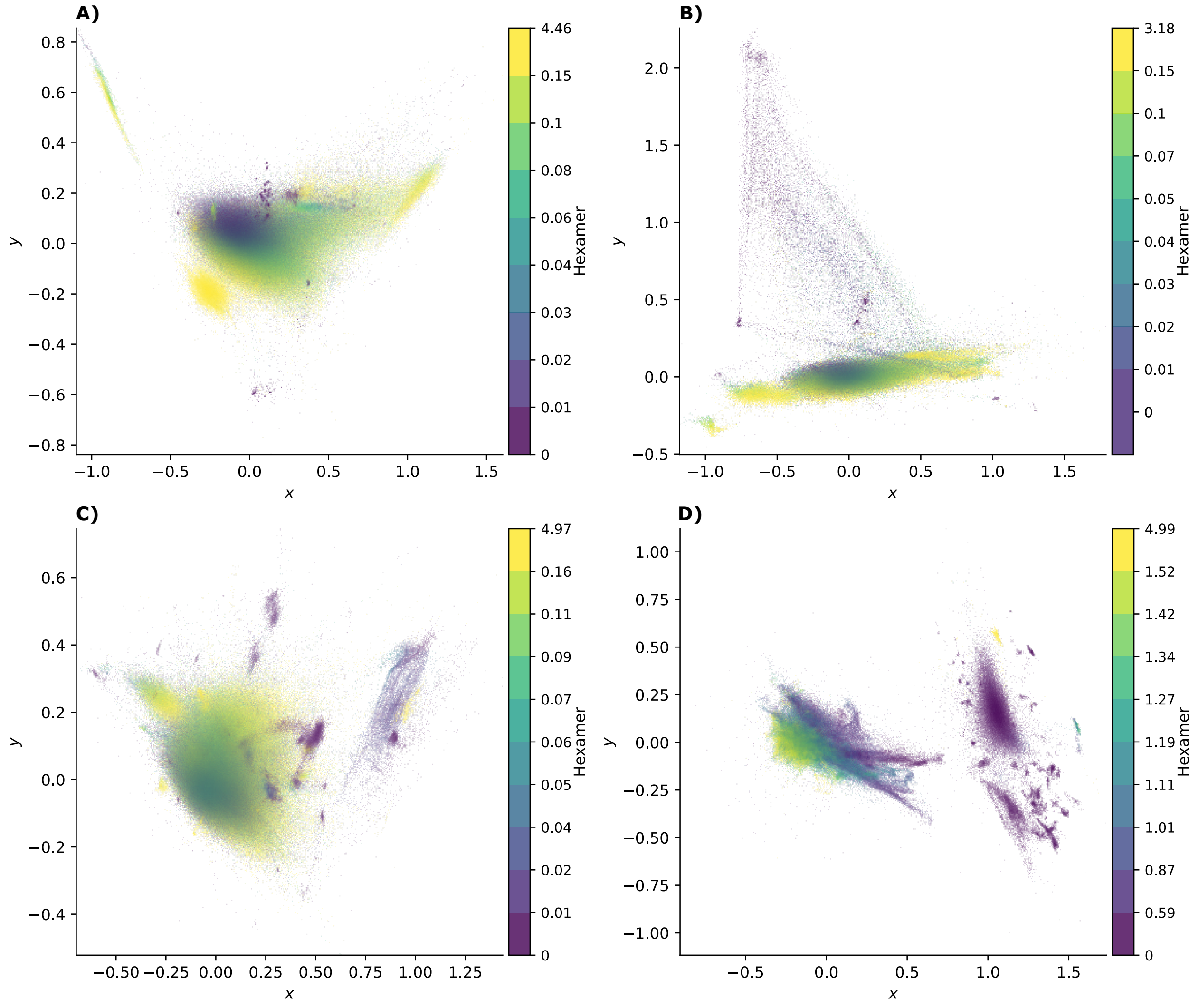


**Supplementary Figure 7.** Read tetranucleotide counts from A) *P. bucephala,* B) *B. lacticolella*, C) *T. albacares*, and D) *B. submarina* projected into two dimensions by PCA, and annotated with estimated coding density. In addition to producing less distinct read clusters than the equivalent latent embeddings from the equivalent VAE, PCA results in the points being less evenly spread across the plot (see Figures S1 and S2). As a result, the *Erwinia* and *Nosema* clusters in *B. lacticolella* are hardly discernible from static plots. The *T. albacares* sample shows a similar pattern, with poor separation between *Kudoa* and host sequences.

#### Interpretation of the latent features

Inspecting the latent dimensions revealed that, for all examples illustrated here, GC strongly correlated with at least one latent dimension. Therefore, GC is one feature that contributes to separating sequences from different components of the sample, though it is not sufficient. Where the distribution of GC content overlaps between the target and cobiont reads, as is the case with the buff tip moth and *Wolbachia*, additional information is needed.

| Species | Correlation with GC | |
|---|---|---|
| | $\mu_0$ | $\mu_1$ |
| *Phalera bucephala* | -0.1 | -0.98 |
| *Blastobasis lacticolella* | 0.99 | -0.01 |
| *Thunnus albacares* | 0.06 | 0.94 |
| *Brachiomonas submarina* | -0.95 | -0.18 |

**Supplementary Table 1.** Correlations between GC content and encoder outputs for two latent dimensions, measured by Spearman's rho. $P < 0.0001$ for all correlations. Both dimensions correlating with GC to some extent may be a consequence of $\beta << 1$.

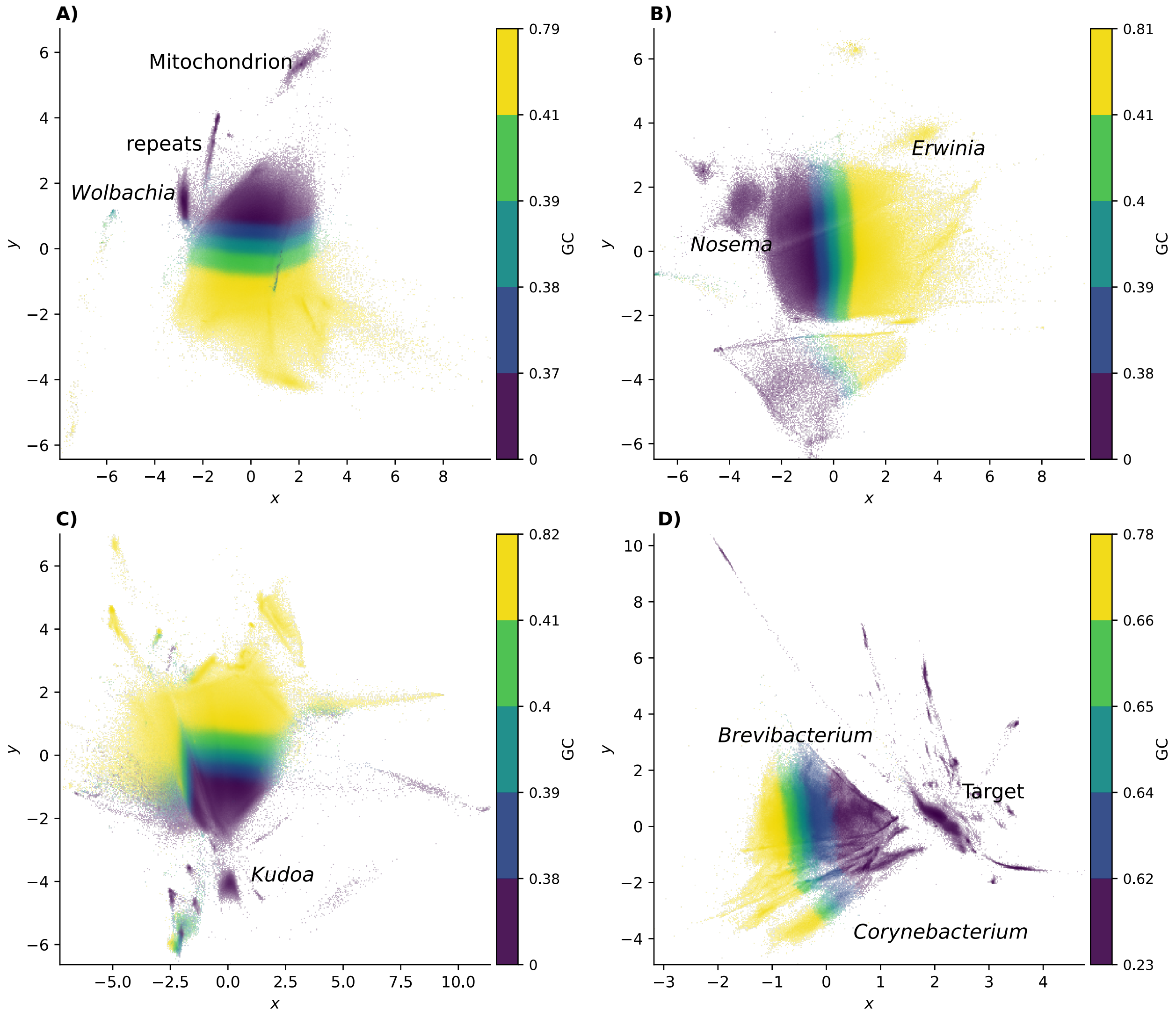


**Supplementary Figure 8.** Projected read tetranucleotide counts for *P. bucephala, B. lacticolella*, *T. albacares*, and *B. submarina* samples (A-D) labelled by GC content quantile. Given the small range of values for intermediate GC contents, only five bins are shown.

#### Impact of extreme base composition on Mash Screen hits

For many lepidopteran samples, Mash Screen returned spurious hits to *Buchnera aphidicola*, *Candidatus Carsonella ruddii*, *Plasmodium reichenowi*, *Plasmodium gaboni*, *Candidatus Sulcia muelleri*, *Candidatus Phytoplasma oryzae*, *Candidatus Phytoplasma mali*, and *Ichthyophthirius multifiliis*

The case of *Carsonella*, which has a GC content of 14%, illustrates the problem of spurious matches due to biased nucleotide content: Sequences corresponding to sketched k-mer hashes with the default $k = 21$ for the *Carsonella* genome included numerous spurious matches to AT-rich regions of insect contigs. The probability of a random match is given by $\frac{1}{(\Sigma^k/g)+1}$, where $g$ is genome size (see https://mash.readthedocs.io/en/latest/sketches.html). Given that the effective size of the nucleotide alphabet $\Sigma$ from which the k-mers are drawn is less than 4 when C and G are rare, the default k-mer length of 21 is expected to be too small to provide sufficient specificity. Indeed, re-sketching the *Carsonella* reference with $k = 31$ using Mash Sketch reduces the number of matches in the reads of the *Chloroclystis v-ata* sample from 169 to 1. Meanwhile 126 of 152 matches to *Wolbachia* remain. However, increasing $k$ may lead to an undesirable loss of sensitivity overall. Devising a strategy to mitigate false positives returned by Mash Screen is beyond the scope of this work, and relevant hits were therefore removed.
